# Behavioral, molecular and neuronal mechanisms involved in recognition memory retrieval under degraded spatial cues in the rat hippocampus

**DOI:** 10.1101/2023.03.14.532607

**Authors:** Magdalena Miranda, Azul Silva, Juan Facundo Morici, Marcos Antonio Coletti, Mariano Belluscio, Pedro Bekinschtein

**Author notes:** Equal contribution and co-corresponding authors. Equal contribution authors (Magdalena Miranda, Azul Silva).

## Abstract

In a constantly changing environment, organisms face the challenge of adapting their behavior by retrieving previous experiences or acquiring new information. Previous research has postulated that this balance between memory generalization and differentiation manifests in a dichotomic manner. When environmental information exceeds a given threshold, activation of a stored representation could initiate retrieval, but below this threshold, a novel event could be encoded with a concomitant remapping of the internal representation in the hippocampus. Here, we examined the hippocampal molecular and neuronal mechanisms underlying retrieval in a cue-degraded environment by combining in vivo electrophysiological recordings and pharmacological manipulations. We developed a memory recognition task that allows a graded decrease in the contextual cues present during retrieval. We found that the manipulation of the number of visual cues was consistent with the activation or not of the contextual memory trace. Retrieval of a specific context memory was reflected by the level of CA3 remapping, demonstrating a clear relationship between remapping and contextual recognition. Also, manipulation of NMDAR activity in the DG-CA3 circuit bidirectionally modulated contextual memory retrieval. The blockade of NMDAR in CA3 impaired recognition in a cue-degraded, but not in a full-cue context, while their activation has the opposite effect. Conversely, blockade of NMDAR in the DG promoted retrieval under an even more cue-degraded environment, while activation had the opposite effect. Our results provide evidence for a flexible interaction between environmental cues and information stored in the hippocampus and give new insights into the biological mechanisms that balance memory encoding and retrieval.

## Introduction

Learned behavior is the consequence of an interaction between the environment and the representations stored in memory. Behavior can depend on whether experiences are novel or similar to previous ones. Since the environment is continuously changing, episodic memory retrieval usually occurs under contextually degraded or modified conditions and the ability to retrieve previous memories despite partial contextual change becomes crucial [1]. However, failure to discriminate between similar memories could be detrimental for episodic memory formation due to interference between stored memories [2-6]. A changing environment forces the brain to make a difficult choice: should we forget about the differences between two similar experiences and retrieve an already stored representation or should we make a new memory? Discarding the distinct details would allow to conceptualize and create rules to predict environmental changes, whereas the creation of a new memory could add important information and new resources for a diverse set of behavioral responses. Under each particular context, the brain should be able to evaluate environmental and internal cues to determine whether that context is familiar or not. Based on the proposed dual role of the hippocampus (HP) in context change and retrieval, computational studies suggested the need for two independent systems [7] to optimize information processing under small or large contextual changes. These studies proposed that unique features of hippocampal subregions could allow the development of computationally distinct and complementary functions needed for correct episodic memory formation and retrieval, known as pattern completion in the case of the CA3 region and pattern separation for the DG region [5, 7-13].

However, many questions remain regarding how context is represented, stored and retrieved by the hippocampus. Hippocampal cells are active as assemblies that can represent space (place cells) [14], time [15, 16], events [17-19] or a conjunctive representation of them [20-22]. Different contexts could be represented as independent maps in the HP [23-25]. Place cells fire specifically when an animal is in a particular location [26], in a way that different place cells encode different positions forming a cerebral spatial map [27-29]. The process that leads to a reorganization of cell ensemble activity when the context changes is known as remapping. Remapping can be complete or partial, depending on the conditions of spatial shifts [30].

But can remapping explain retrieval or encoding of spatial memories that sustain behavior? Despite the intuition of place cell importance for episodic memory, there is still no clear evidence that remapping is directly related to the expression of different contextual memories and there is not even a clear understanding of the precise relationship between contextual changes and remapping. One reason for this is that most studies have only controlled the observable properties of the environment but did not analyze behavioral outputs that could give some insight into the inferences the animal is making about a particular environment or experience. In other words, the variability of circuit activation in the same physical environment has been overlooked, variability that could be influenced by motivation [31, 32], attention [33] and experience [34]. Moreover, it is not clear how this individual mnemonic variability (i.e. behaving as being in a familiar context or not) relates to or influences the internal context representation associated with place cell activity.

Both computational models and theoretical dual process models (DPM) have considered episodic memory retrieval as dichotomic, with reactivation of the neuronal ensemble occurring when the available information exceeds a given threshold [35, 36]. From a place cell perspective, above this threshold a pattern completion process would control the amount of remapping and allow reactivation of the original representation from only degraded or incomplete information and subsequent memory retrieval. Below this threshold a pattern separation process would transform correlated information in independent non-overlapping representations supporting the event to be coded as a novel experience [37]. Experimental evidence suggests a role of DG in pattern separation and of CA3 in pattern completion, both at the electrophysiological and behavioral level [38-41]. In addition, NMDA receptors (NMDARs) in DG and CA3 have been implicated in behavioral memory discrimination and rate remapping [29, 30]. Although DG/CA3 circuit connectivity could be the substrate for pattern separation/pattern completion, synaptic plasticity could represent an adequate functional strategy to regulate the efficiency of this process [42]. There is also evidence to suggest a role of plasticity related mechanisms for the retrieval of engram memory representations under a cue degraded context. Previous studies showed that NMDARs are particularly relevant to recover memory from a cue degraded context [43] and to allow CA1 neurons to maintain place field characteristics between these conditions [44, 45]. Since some theories suggested that the DG-CA3 circuit of the HP mediates the dynamic competition between the process of differentiation (supported by pattern separation) and generalization (supported by pattern completion), understanding the nature of this interaction could disentangle which kind of relationship explain the nature of context representation based episodic memory.

In this work, we study the interaction between pattern separation and pattern completion during episodic like-memory at the behavioral (i.e. memory retrieval) and electrophysiological (i.e. remapping) level. To model episodic memory retrieval in an incidental context without the bias of motivational influences over contextual representations, we developed a context-degraded task. Taking into account the fundamental role of the balance between these processes for episodic memory function, we decided to test the role of molecular mechanisms as potential modulators of this balance in the DG-CA3 circuitry and analyze CA3 neuronal activity. We found that the internal context representation, seen as the efficacy of memory retrieval to guide behavior, could be explained by the context representation by CA3 neurons. Furthermore, we showed that memory formation and its retrieval can be modulated by altering the activity of NMDARs in the DG-CA3 circuitry.

## Methods

### Subjects

For the 2-days version experiments, subjects were 197 male Wistar rats that weighed approximately 250-300 g at the beginning of the study. Rats were housed in groups of two to five. Rats were food deprived to 85-90% of their free feeding weight to increase spontaneous exploration, except during recovery from surgery, where food was available *ad libitum*. Water remained available *ad libitum* throughout the study. These studies were conducted in accordance with the National Animal Care and Use Committee of the University of Buenos Aires (CICUAL) and the University of Favaloro (CICUAL UF 2021-006)

For the 1-day version experiments, data was obtained from 10 male Wistar rats that approximately weighed 300–400g. All 10 rats were used for the behavioral analysis (Fig 1); 4 rats were implanted for neuronal recordings (Fig.3). Rats were housed separately in transparent plastic cages. Rats were water deprived to 85-90% of their free drinking weight to increase spontaneous exploration, except during recovery from surgery, where water was available *ad libitum*. Food remained available *ad libitum* throughout the study and water was available for 20 minutes each day. These studies were conducted in accordance with the National Animal Care and Use Committee of the University of Buenos Aires (* COPDI-2021-02755266-UBA-DDG%FMED).

**Fig 1.**
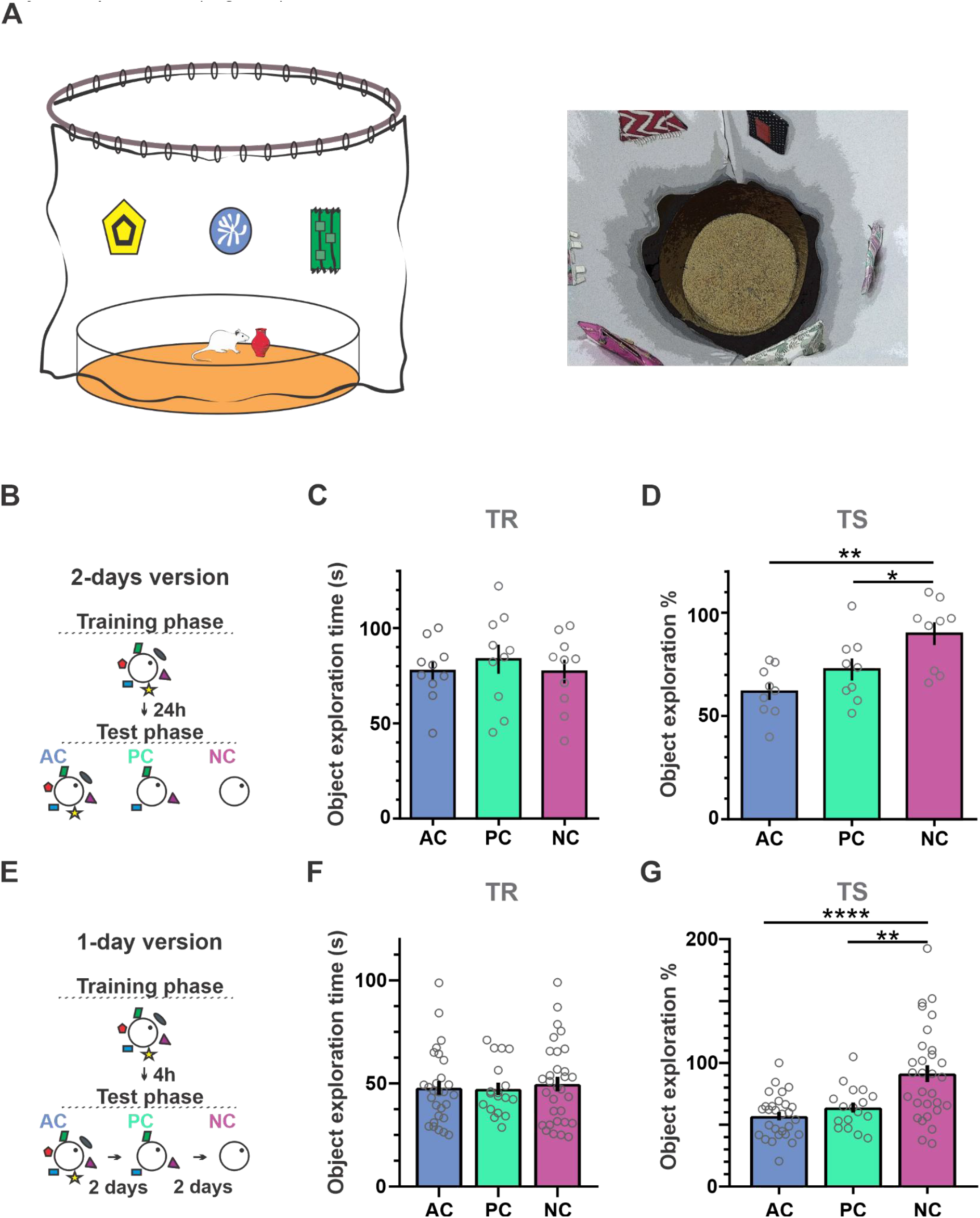
The Associative Retrieval Task. **(A)** AR task. “All cue” (AC, blue), “Partial cue” (PC, green) and “No cue” (NC, violet) conditions. **(B)** Schematic illustration of the 2-day version of the task. **(C)** Total object exploration time during the training session for AC, PC and NC during the 2-day version of the task. Rats spent an equal amount of time exploring the object during the training phase under the AC, PC and NC conditions. One Way RM ANOVA F=0.74, p=.453, n=9. **(D)** Percentage of object exploration in the presence of a variable number of cues (AC, NC and PC) in the test session of the 2-day version of the task with respect to training. RM One way ANOVA, F=8.27, p=.006; One sample t test against 100% AC t=9.39, p<.0001; PC t=5.18, p=.0008; NC t=1.83, p=.105. **(E)** Schematic illustration of the 1-day version of the task. **(F)** Total object exploration time during the training session for AC, PC and NC during the 1-day version of the task. Two way RM ANOVA, interaction: F =.5755, p = .7484, n=10, sessions = 75, each dot represents a session. **(G)** Percentage of object exploration in the presence of a variable number of cues (AC, NC and PC) in the test session of the 1-day version of the task with respect to training. Two way RM ANOVA, main effect condition F = 16.04, p < .0001.Tukey’s post hoc test: **** p<.0001 AC vs NC; p=.25 AC vs PC ; ** p =.001 NC vs PC. One sample t test against 100% AC t =14.45, p<.0001; PC t=8.76, p<.0001; NC t = 0.37 p=.91.

For both versions of the experiment, rats were housed on a reversed 12 h light/12 h dark cycle (lights on 19:00-07:00) and all behavioral testing was conducted during the dark phase of the cycle.

### Surgery and Cannulation

Rats were bilaterally implanted in the DG, Prh or CA1 region with 22-gauge indwelling guide cannulas. Subjects were anesthetized with ketamine (Holliday, 74 mg kg-1, ip) and xylazine (Konig, 7,4 mg kg-1, ip) and were placed in a stereotaxic frame (David Kopf Instruments, Tujunga, CA) with the incisor bar set at -3.2 mm. Guide cannulas were implanted according to the following coordinates, measured relative to the skull at bregma [46]: Prh AP-5,5 mm, LL ± 6,6 mm, DV -7,1 mm; DG AP -3.9 mm, LL ± 1.9 mm, DV -3.0 mm; CA3 AP -3.6 mm, LL ± 3.6 mm, DV -3.6 mm. The lateral separation of 1.7 mm between DG and CA3 coordinates is sufficient to allow a distinction between the actions of these infusions since previous reports indicate that the spread of an infusion of 0.5 μL volume is approximately a 1.0 mm diameter sphere that is consistent for up to an hour after the infusion [47, 48].

The cannulas were secured to the skull using dental acrylic and three jeweler screws. Obturators, cut to sit flush with the tip of the guide cannulas and with an outer diameter of 0.36 mm, were inserted into the guides and remained there except during infusions. At the completion of each surgery, an antibiotic was applied for three days (Enrofloxacin; 0.27 mg kg-1, Vetanco, Arg). Animals were given at least 7 days to recover prior to drug infusions and behavioral testing.

### Surgery and tetrode implantation

Rats were implanted with a custom-designed and 3-D printed microdrive with sixteen independently movable tetrodes, eight tetrodes aiming at CA3 and 8 tetrodes aiming at CA1 (data not shown). Subjects were anesthetized with isoflurane (air flow: 0.8–1.0 l/min, 4% isoflurane for 3 minutes for induction, 0.5%–3% isoflurane for surgery, adjusted according to physiological condition). In addition, they received subcutaneous injections of dexamethasone and a local injection of lidocaine at the start of the surgery. Isoflurane was gradually reduced from 3% to 0.5%. Upon induction of anesthesia, the animal was fixed in a stereotaxic frame (David Kopf Instruments, Tujunga, CA). Temperature was controlled with a servo-controlled heating pad (37°C ± 0.5; Fine Science Tools, Vancouver, Canada). Craniotomies for tetrode implantation were drilled in the skull above the CA3 (AP: -3.6; ML:±4, DV: -3.4) and CA1 region and tetrodes were then implanted. Tetrodes were made of four twisted 12 μm tungsten wires (CFW, USA). Electrode tips were gold-plated to reduce electrode impedances to around ∼200 kΩ at 1 kHz and dyed with Dil (Thermo-Fisher, United States) to check tetrodes positions. During surgery, tetrode tips were lowered to 1 mm above the structure. After 5 d of recovery, tetrodes were moved gradually until they reached CA3. Neuronal spiking activity and LFP were recorded daily in the home cage and every two days in the behavioral task. Units recorded in different sessions were considered independent.

### Infusion procedure

Depending on the experiment, rats received bilateral infusions of Emetine (50 μg μl^-1^/ 0.5 μl side; Sigma-Aldrich), D-Cyc (D-4-amino-3-isoxazolidone, 20 μg/μl; Sigma-Aldrich), AP5 (2-amino-5-phosphonopentanoate, 2 μg μl-1/ 0.5 μl side), AMPA/kainate DNQX (6,7-dinitroquinoxaline-2,3-dione, 1 μg μl^-1^/ 0.5 μl side; Sigma-Aldrich), Nimodipine (7.5 μg/ 0.5μl, Tocris) or Vehicle (Veh, saline/ DMSO 5% for DNQX/ DMSO 2%-Tween 2% for ANA-12) at different times during the behavioral task. Bilateral infusions were conducted simultaneously using two 5-μl Hamilton syringes that were connected to the infusion cannulas by propylene tubing. Syringes were driven by a Harvard Apparatus precision syringe pump, which delivered 0.5 μl to each hemisphere over 2 min. The infusion cannulas were left in place for an additional minute to allow for diffusion. At least 3 days were allowed for washout between repeated infusions. One or two days after the behavioral procedure, all animals were infused with 0.5 μl of 5% methyl blue solution through the cannula and were sacrificed 15 minutes later. Cannula localization was verified and was correct in over 95% of the surgeries. Only the behavioral data for animals with correctly implanted cannulae were included in the analysis.

### Recording procedure

During the recording sessions, neurophysiological signals were acquired continuously at 20 kHz on a 256-channel Amplipex system (http://www.amplipex.com). The microdrive had three small leds attached to track the rat’s position on the open field that were recorded by a digital video camera at 30 frames/s. LED locations were detected and recorded online with the same software used for the neurophysiological signals.

### Behavioral procedure

For the associative retrieval (AR) task, we used a circular open field (90 cm diameter x 45 cm high) made of black plastic surrounded by six spatial cues placed over a curtain that enclosed the context preventing visibility of other distal cues in the room. All six cues were detachable from the curtain to modify their number in different testing conditions. All cues consisted of carton and cloth shapes with approximately the same surface (850 cm^2^). The open field was situated in the middle of a dimly lit room. The open field’s floor was covered with wood shavings only for pharmacological experiments. A video camera was placed over the arena and both sample and choice phases were recorded for later analysis.

For the 2-days version of the AR task, each rat was handled for 2 days and then habituated to the arena with the corresponding cues for 10 min a day for 5 days before exposure to the objects. After habituation, during the training phase, animals were exposed for 10 min to an object in a pseudorandomized position in the arena in presence of 6 cues. The test phase lasted 10 min and was made 24 hours after the training phase. During this phase, an identical copy of the object was placed in the same position as during training but in the presence of a variable number of distal contextual cues. This was done in order to create an object in context memory. For the ‘all cue’ condition (AC), object presentation occurred in the presence of all the cues presented during training. In the ‘partial cue’ condition (PC), object presentation was done in a context with half of the cues (3), and in the ‘two partial cue’ condition (PC2) only 2 contextual cues were present. Finally, for the ‘no cue’ (NC) condition the object was presented in the absence of spatial cues (see Fig.1).

For the 1-day-version of the AR task, the procedure was similar except that only during habituation small drops of water were spread in the open field to increase context exploration. Animals were habituated to the PC and SC condition. Moreover, the duration of training and test phase was 13 minutes to increase the amount of signal recorded.

For the reconsolidation experiments, rats were habituated in both the open field for 5 days and a triangular arena for 3 days. After habituation, animals were trained in the AR task for 10 min and, 24 hours later, a retrieval session was performed. During this session (TS1), animals were exposed to an empty context for 10 min with a variable number of contextual cues (depending on the experiment). During the test session (TS2) 24 hours after, a copy of the original object was presented next to a novel object in the triangular arena for 5 min. For the experiment on Fig. 2C, during the first day animals were trained in the AC context with an object A and in the NC condition with the object B. These two training sessions were given in a pseudorandom order and separated by 2 h. In the evaluation, a group was exposed to an empty AC context with all the cues and another group was exposed to the empty AC context with no cues. During a second evaluation, memory for the presented objects (A or B) was tested against a novel object (C or D) in the triangular arena for 5 min. The order of the two test sessions was counterbalanced.

**Fig. 2.**
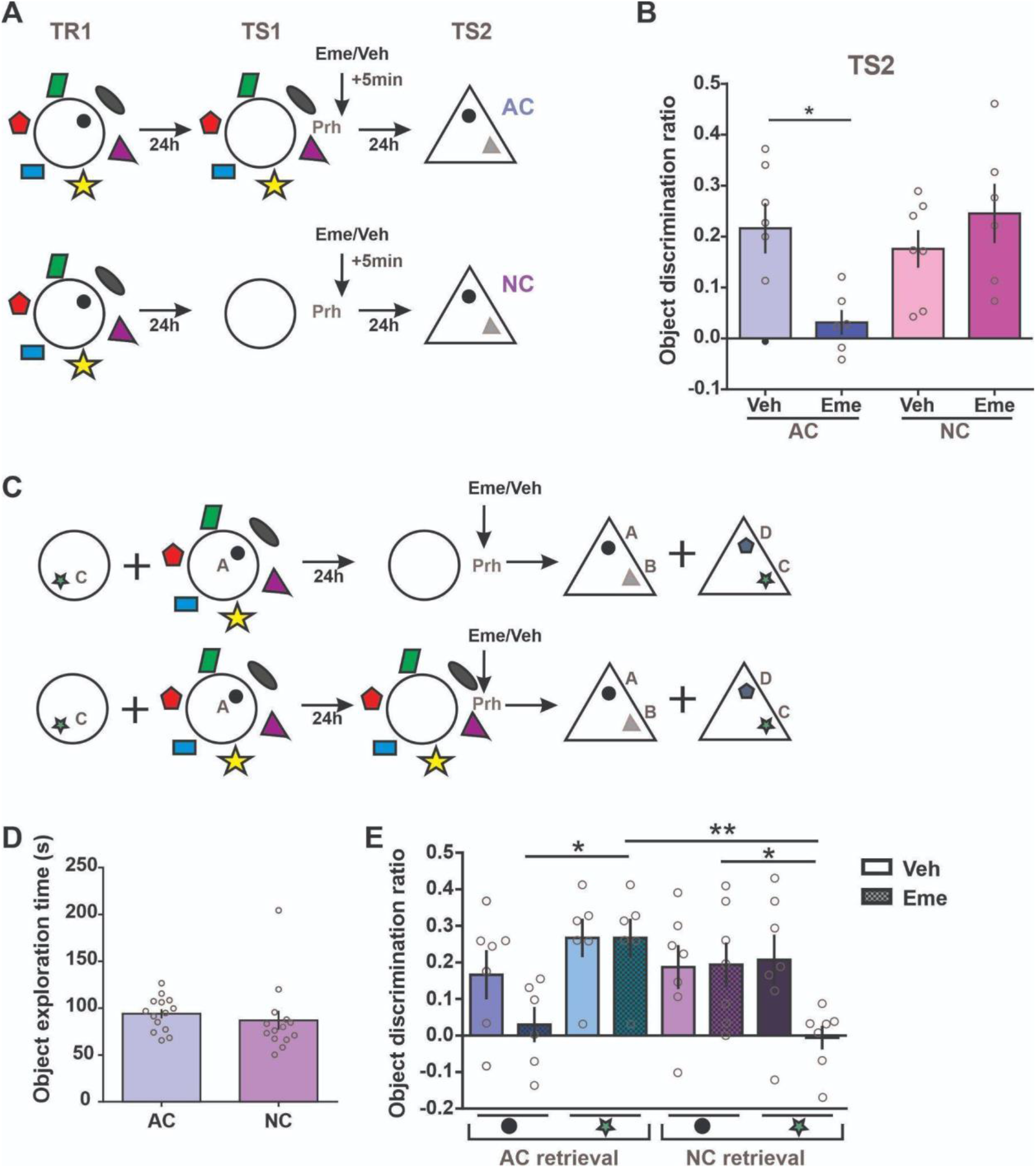
Contextual cues are used to guide retrieval of the relevant object memory trace in the Prh. **(A)** Procedure and time points of infusions. Animals were trained in the AR task and 24h after were exposed to an empty AC or NC context (TS1) and immediately after they received an Emetine (Eme) or Vehicle (Veh) infusion in the Prh. A test session was given to evaluate the original object against a novel one (TS2). **(B)** Discrimination ratio for the TS2 test session, 24h after exposure to an empty context with AC or NC followed by an Emetine (dark) or Veh (light) infusion. RM Two way ANOVA: F_interaction_=6.25, p=.029, F_condition_=2.90, p=.117, F_Drug_=2.50, p=.142, n=6-7. One sample t test against 0: NC-Veh t=4.77 p=.003, NC-Eme t=4.22 p=.008, AC-Veh t=4.38 p=.005, AC-Eme t=1.30 p=.252. **(C)** Experimental protocol and time points of the Emetine or Vehicle infusion. Animals were trained to an object A (circle) in the NC context and an object C (star) in the NC context. The next day they received an exposure session to an empty AC or NC context (no object) and immediately afterwards they were infused with either Emetine (dark) or Vehicle (light) in the Prh. Lastly, object memories were tested against novel objects in a different context 24h after (TS2). **(D)** Object exploration time during training in the AC or NC context. T-test transformed data t=1.66 p=.121 n=6-7. **(E)** Discrimination ratio for the A or C objects against a novel object during TS2, 24h after exposure to an empty AC or NC context followed by Emetine (gridded) or Vehicle (smooth) in the Prh. Two Way ANOVA: F_interaction_=2.08, p=.132, F_condition_=4.05, p=.020, F_Drug_=4.77, p=.039, n=6-7. * p <.05, ** p <.01. Individual values used to calculate the mean and SEM are presented as dots.

### Data Analysis

#### Experimental design and statistical analysis

In every experiment, pharmacological infusions or experimental conditions were counterbalanced between trials. For the AR experiments, results were presented as the Object exploration %, ie. percentage of exploration during the test phase with respect to the training phase ((t^test^/t^training^)* 100). For reconsolidation experiments, results during the TS2 session were expressed as a discrimination ratio that was calculated as the time exploring the novel object minus the time exploring the familiar object divided by total exploration time ((t^novel^-t^familiar^)/t^total^). Exploration of an object was defined when the rat had its nose directed towards an object at a distance of 2 cm or less, or touching the object with its nose. Leaning on the object with the head facing up does not count as exploration. Climbing or sitting on the object is not included as exploration. For rearing quantification, a rearing was defined as the animal climbing into an upright position, resting with hind limbs either in the air or leaning against a wall or object (and only when the head was oriented up but not exploring the object). For all quantifications, scoring was performed by two different experimenters blind to the phase of the task and the experimental condition.

One sample t tests were used to compare the percentage of exploration with respect to 100%, as a measure of memory. Additionally, paired t tests were used to compare between conditions. In the cases where data did not meet normality criteria, Wilcoxon signed-rank test was used instead. For experiment shown in Fig. 1 B,C,D, the two days version of the AR task, animals were tested three times: during the first trial a third of the animals were tested in the AC condition, a third in the PC and a third in the NC; during the second trial, half of each third was tested in one of the conditions and half in another; and for the third trial the remaining condition was tested. Animals whose percentage of exploration exceeded 2SD from the group mean were excluded from the analysis. Behavioral response was analyzed using a Repeated Measures (RM) ANOVA. Data that did not meet normality criteria was transformed before analysis.

For experiments in Fig 1 E,F and G, the 1-day version of the AR task, animals were tested every two days in one condition of the three conditions in a pseudo randomized order. Behavioral response was analyzed using a Repeated Measures (RM) ANOVA (Fig S.1). Neuronal response in Fig 3 B and C was analyzed with a Kruskal-Wallis test and in Fig 3 E and F a general mixed model was used with rats as a random factor. The best general linear model was selected using an ANOVA comparing models with increasing complexity. For experiments in Fig. 2, 4B, 5B, C, E animals were tested twice. During the first trial, half the animals received an infusion of Eme/AP5/D-Cyc (depending on the experiment) and the other half received an infusion of the corresponding Vehicle of each drug. In the second trial, animals received an infusion of Eme/AP5/D-Cyc or the corresponding Vehicle according to what they received in the first trial. For the training phase, the percentage of exploration time for each object was compared using a RM ANOVA. And for the case of Fig. 5F, animals were tested three times with either Veh/Veh, Veh/Eme or Nimo/Eme. In Fig. 4D animals were tested in two opportunities. In the first trial half the animals received a Vehicle infusion in the Prh and a Vehicle or AP5 in the CA3/GD, while the other half was infused with Emetine in the Prh and Vehicle or AP5 in the CA3/GD. In the second trial, animals that had received Vehicle in the Prh now had an Emetine infusion and vice versa.

**Fig 3.**
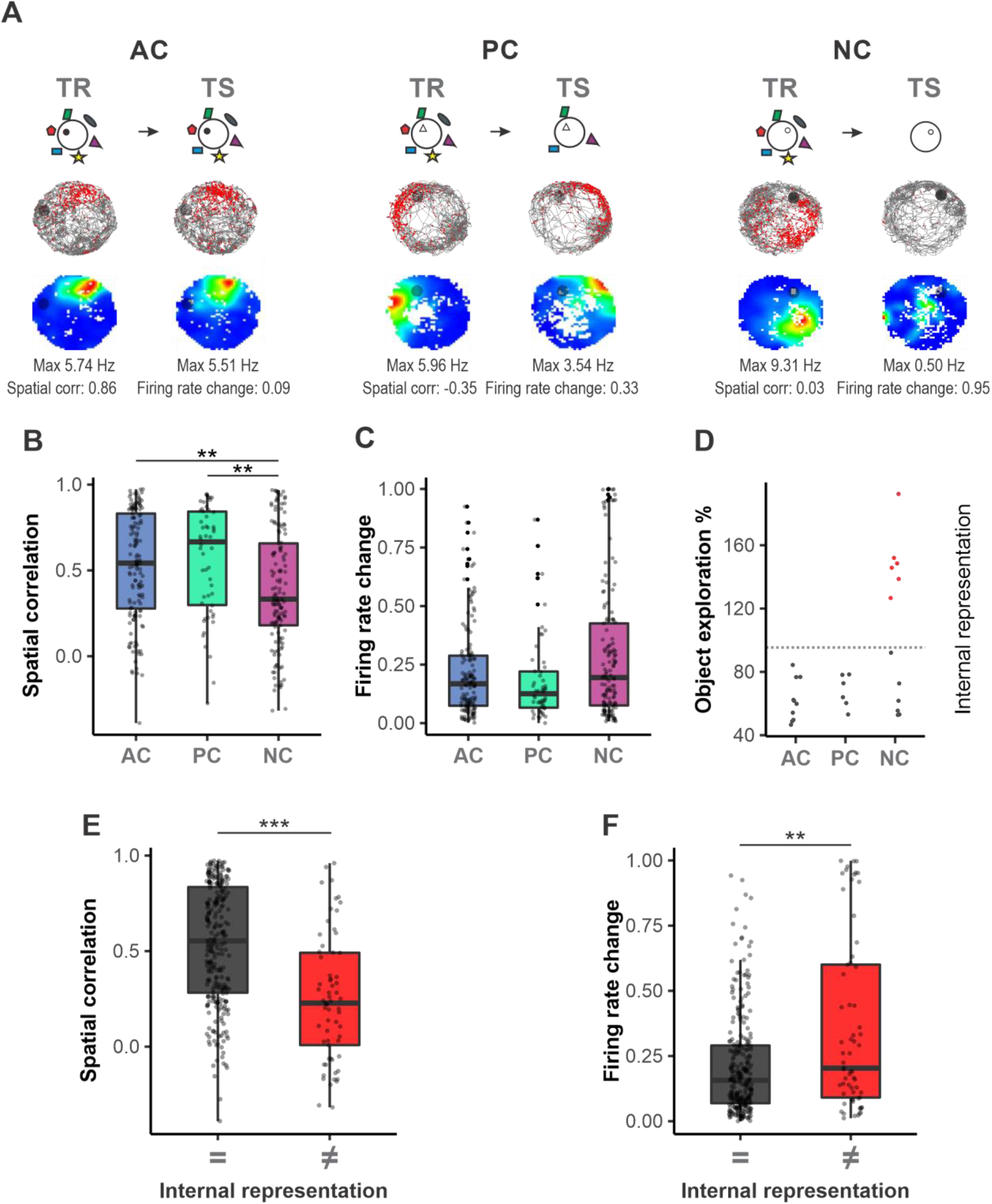
CA3 place cells remapping correlates with memory recall. **(A)** Different types of ca3 place cell responses. Place maps for training session (left) and test session (right). Spatial correlation (Pearson correlation between the firing rate in both phases) and firing rate change (division between the subtraction of training firing rate and test firing rate and their sum) were calculated for each neuron. Upper panel: gray lines represent the animal’s trajectory, red dots, the location of each action potential, and the black circle, the object. Lower panel: color-coded firing maps normalized by the min and the max firing rate of each neuron. **(B)** CA3 spatial correlation sorted by condition. Only in NC sessions, place cells had a significantly lower spatial correlation. Moreover, there were no differences between the AC and PC conditions. Although half of the cues were removed in PC, there were no differences in neuronal activity between PC and AC. n=332, Kruskal-Wallis test NC-AC p= .0027, NC-CP p= .0074 and AC-CP p= .8503. **(C)** CA3 firing rate change sorted by condition. There are no differences between conditions. n=332, Kruskal-Wallis test NC-AC p= .3430, NC-CP p= .0720 and AC-CP p= .4788 **(D)** Object exploration percentage for the three conditions and for the internal representation: sessions were divided by a context discrimination threshold (mean (OE% AC OE% PC) + 2 STD) red dots represent sessions where animals discriminated between contexts and gray dots, sessions where animals didn’t discriminate. **(E)** CA3 spatial correlation sorted by internal representation. n=332, LMM p =2.311e-06 **(F)** CA3 firing rate change sorted by internal representation. n=332, LMM p = .006

**Fig. 4.**
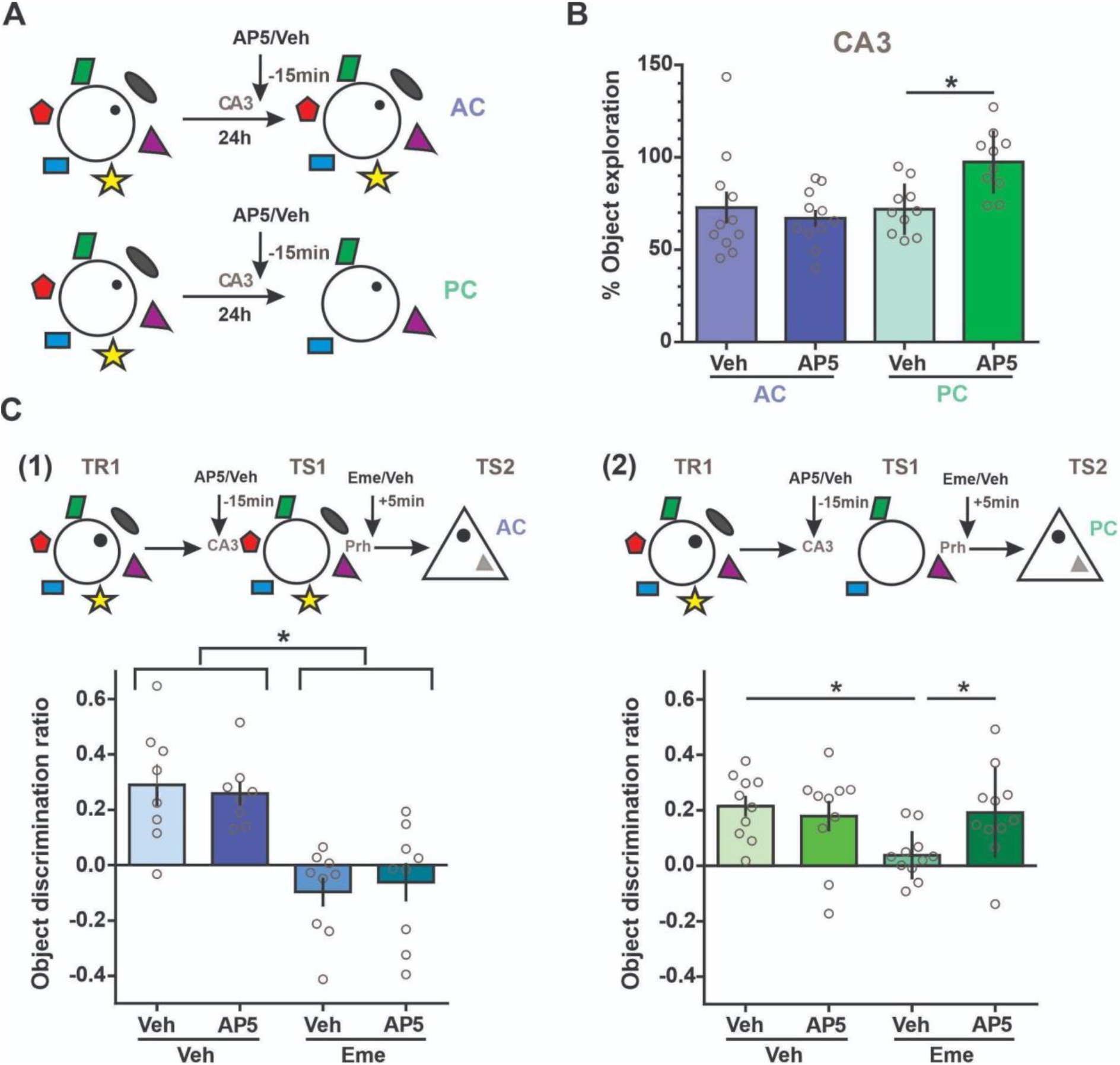
NMDAR requirement for associative retrieval of object in context memory in a cue degraded context. **(A)** Time of infusion of AP5 or Vehicle (Veh) in the CA3 region of the HP 15 min before the test session in the AC and PC conditions of the task. **(B)** Percentage of exploration for the object presented in the presence of the AC or PC conditions during the test session, in animals infused with Vehicle (light) or AP5 (dark) in the CA3 region of the HP. Two Way ANOVA: Logaritmic transformation n=10-11, F_interaction_=4.50, p=.047, PC_Veh-AP5_ p= .042. Wilcoxon test against 100%, AC-Veh W=-48, p=.032; PC-Veh W=-66, p=.001; AC-AP5 W=-55, p=.002; PC-AP5 t=-7, p=.769. **(C)** Experimental protocol used for the task. Animals were trained in the AR task and 24h after they received a Veh or AP5 infusion in CA3 and 15 min after were exposed to an AC (1) or PC (2) context, and immediately after received an infusion of Vehicle or Emetine in the Prh. In a final test session 24h after, memory for the original object was tested in another familiar context against a novel object (TS2). **(D)** (1) Discrimination ratio for the TS2 session of the task, 24h after exposure to an empty original context followed by an Emetine or Vehicle infusion in animals that had previously received AP5 or Vehicle infusions. Two way ANOVA: F_interaction_=0.26 p=.618, F_Veh-Eme_=35.93 p<.0001, F_Veh-AP5_=0.001 p=.970, n=8-9. One Sample t test against 0, Veh-Veh t=3.81 p=.007, Veh-Veh t=3.81 p=.007, AP5-Veh t=5.98 p=.0006, Eme-Veh t=1.85p=.101; Eme-AP5 t=0.89 p=.3996. (2) Discrimination ratio for the TS2 session of the task, 24h after exposure to an empty partial cue context followed by an Emetine or Vehicle infusion in animals that had previously received AP5 or Vehicle infusions. Two way ANOVA: F_interaction_=4.42 p=.049, n=10-11. One sample t test against 0: Veh-Veh t=5.83p=.0003, Veh-Veh t=3.23p=.010, AP5-Veh t=1.44p=.181, Eme-Veh t=3.90 p=.0030; Eme-AP5 t=0.89 p=.40. * p <.05. Individual values used to calculate the mean and SEM are presented as dots.

**Fig. 5.**
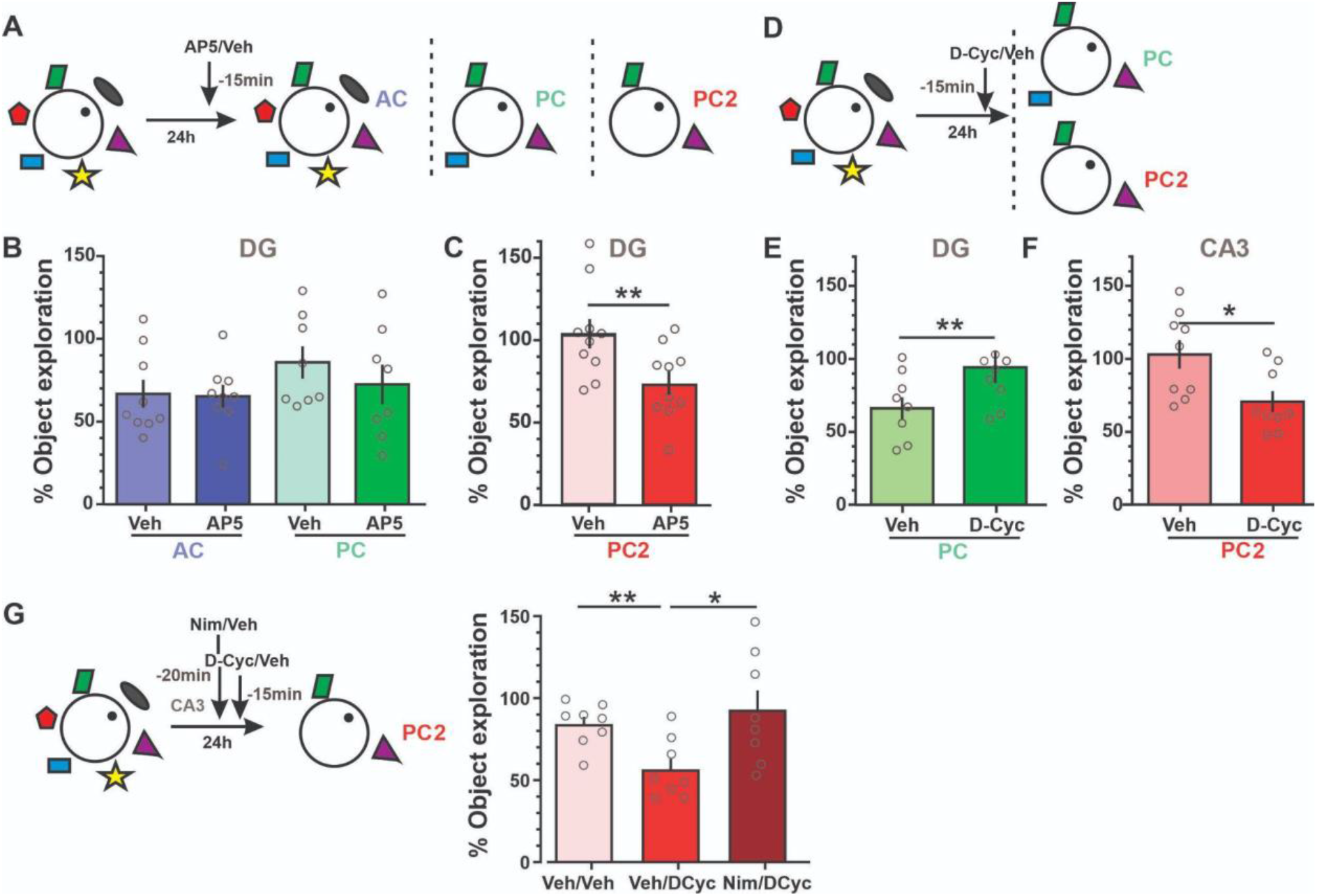
NMDAR agonist in the CA3 increases retrieval capacity in the presence of partial cues, in a L-type voltage gated calcium channel dependent manner, while in the DG region has an opposite effect. **(A)** Time of infusion of the NMDAR agonist D-Cyc or Vehicle (Veh) in the DG of the HP 15 minutes before the evaluation session in the AC, PC or PC2 condition of the task. **(B)** Percentage of exploration for objects presented during the test session under the AC or PC conditions of the task in animals infused with either Vehicle (light) or AP5 (dark) in the DG. Two Way ANOVA GD: n=8-9, F_interaction_=0.42, p=.527, F_condition_=1.93, p=.185, F_drug_=0.65, p=.432. One sample t test against 100%, Veh AC t=3.94, p=.043; AP5 AC t=5.05, p=.001; Veh PC t=1.467, p=.186; AP5 t=2.30, p=.055. **(C)** Percentage of exploration in the PC2 context during the test session in Vehicle infused (light) or AP5 infused animals (dark). Paired t test t=3.26, p=.010, n=10. One sample t test against 100%, Veh p=.669 t=0.44, AP5 p=.005 t=3.65. **(D)** Time of infusion of D-Cyc or Vehicle in the DG or CA3 region of the HP 15 min before the test session in the PC or PC2 condition. **(E)** Percentage of object exploration during the test session in the PC condition of the task in animals infused with Veh (light) or D-Cyc (dark) in the DG previous to the session. Paired t test: t=4.198 p=.004, n=8. One sample t test against 100%: Veh t=4.52 p=.002; D-Cyc t=0.54 p=.601). **(F)** Percentage of exploration during the test session in the PC2 condition of the task after animals were infused with Veh (light) or D-Cyc (dark) in the CA3 region. Paired t test: t=3.03 p=.016, n=9. One sample t test against 100%, Veh t=0.31, p=.767; D-Cyc t=4.15, p=.003). **(G)** (Left) Time of infusion of Nimodipine (Nimo)/Veh and D-Cyc/Vehicle in the CA3 region of the HP 20 and 15 min respectively before the test session. (Right) Percentage of exploration during the test session in the PC2 condition of the task after animals were infused with Veh/Veh (light) or Veh/D-Cyc (red) or Nim/D-Cyc (dark) in the CA3 region. RM One way ANOVA: F=5.44, p=.037, n=8. * p <.05, ** p <.01. Veh/Veh t=3.62 p=.0085, Veh/D-Cyc t=6.717 p=.0003, Nimo/D-Cyc t=0.60 p=.5667. Individual values used to calculate the mean and SEM are presented as dots.

#### Putative internal representation of the context

In order to separate sessions of the 1-day version of the task in terms of the animal’s dichotomic behavioral output (sessions where animals discriminated between contexts from sessions where they did not), we used the distribution of the behavioral output variable (percentage of object exploration) from the two session groups that showed memory retrieval (AC and PC). Sessions with an object exploration rate < 2SD of the mean were considered as sessions where animals recognized the context as the same one experienced in TR (retrieval sessions). Sessions with an object exploration rate > 2-SD were classified as sessions where rats behaved as if the context was different from the experienced in the TR (non-retrieval). For the analysis of the categorical data in the 1-day version, we performed a general mixed model with rats as a random factor. In a similar manner, for the 2-day version of the task, we used the pooled distribution of the AC and PC to determine the +2SD threshold for memory retrieval. To analyze the categorical data in the 2-day version, we performed MacNemar’s Chi-square tests considering whether animals performed above or below 2SD in the Veh condition and whether they changed their performance with the different drug infusions. Experiments were pooled according to the effect of each drug. The “memory boosting” group includes the experiments of D-Cyc in CA3 and AP5 on the DG (both on the PC2 condition) and the “memory impairing” group includes the experiments of the D-Cyc on the DG and the AP5 on CA3 (both in the PC condition).

#### Spike sorting and place cells

Neurophysiological and behavioral data were explored using NeuroScope http://neurosuite.sourceforge.net) [49]. Spikes were sorted in two steps, first automatically, using a custom made software, KlustaKwik (http://klustakwik.sourceforge.net)[50], followed by manual adjustment of the clusters (using Klusters software package; http://klusters.sourceforge.net) [49]. Only units with clear refractory periods and well-defined cluster boundaries were taken into account for the analysis (Harris et al., 2000). In order to be included in the analysis, only units with a stable recording during the whole session were considered. Briefly, animals were recorded in their home cage during 10 minutes after the training phase and after the test phase until they fell asleep. if a unit wasn’t active in one of the visits to the open field, it must be active in the home cage to be included in the analysis. We recorded a total of 1,307 well-isolated units from CA3 of 3 freely moving rats in 26 sessions. Of these, 326 were considered putative place cells. Units were considered place cells if they fulfill 3 criteria: a) their spatial information content [51] was higher than 0,15 bit/spike b) the information content of the unit was different to chance level (computed by shuffling the spike train of the cell and rat position 100 times, calculating the information content each time and establishing the quantile 99% as threshold) in at least one of the phases of the recording day and c) their mean firing rate was higher than 0.1 Hz in at least one of the phases of the recording day.

#### Place cell activity

Only data recorded during epochs when the rat was moving faster than 10 cm/s were used (exploratory epochs). Spiking data and animal’s position were sorted into 5-cm × 5-cm bins to create raw maps of spike counts and occupancy. A Gaussian kernel filter (s.d. = 9 cm) was applied to both maps. A smoothed rate map was constructed by dividing the smoothed spike map by the smoothed occupancy map. The smoothed rate maps of the training and test phase were used to compute the mean and peak firing rates in the open field as well as the spatial correlation and firing rate remapping.

#### Spatial correlation and firing rate change

In this analysis, bins visited less than 15 ms and less than 4 times were excluded to avoid artifacts. Moreover, only bins visited in both test and training phases were included. For the spatial correlation measure, smooth rate maps of individual place cells in the training and test phase were compared with a bin-by-bin Pearson′s correlation. Firing rate change between the training phase and the test phase was estimated for each cell by dividing the difference between the mean firing rate of the training and testing phase, and their sum. For the Bootstrap sequential LRT for the number of mixture components, we used an equal variance model with 999 replications.

### Histology

At the completion of behavioral testing, rats were anesthetized by IP injection with 2 ml of Euthatal (Rhône Merieux) and perfused transcardially with phosphate buffered saline (PBS), followed by 10% neutral buffered formalin. The brains were removed and postfixed in formalin for at least 24 hours before being immersed in 20% sucrose solution until they sank.

To verify the site of the infusion, sixty-μm sections of the brain were cut on a freezing microtome encompassing the extent of the injector track. Every fifth section was mounted on a gelatin-coated glass slide and stained with cresyl violet. Slides were examined under a light microscope to verify the location of the injections (Fig. S2A-D,G).

To verify the tetrode’s position, seventy-μm sections of the brain were cut on a freezing microtome. Each section of a relevant part of the hippocampus was mounted on a gelatin-coated glass slide and was examined under a red fluorescent microscope. All tetrodes of the 8-tetrode bundle were identified by finding the tip of each electrode across sections (Fig. S2E,F).

## Results

### The associative retrieval task as a measure of the ability to retrieve original experiences under a cue degraded condition

To determine whether rats would be able to retrieve experiences under degraded environmental cues, we designed an adapted version of the spontaneous object recognition task that we named the associative retrieval (AR) task [52-54]. Rodent inherent preference for novelty requires engaging retrieval to determine an event as familiar or novel. During a training session (TR), animals incidentally associated a novel object with a context that included 6 distal cues. They were later tested (TS) with an identical copy of the object in the same location but under a variable number of the original contextual cues: the “all cues” (AC), the “partial cues” (PC, only 3 distal cues) and the “no cues” (NC, no distal cues) conditions (Fig. 1A, 1B, 1E). For the pharmacological experiments, animals were tested 24 h after the training session (2-days-version) and for the electrophysiological experiments, animals were tested 4 h after the training session (1-day-version). The time animals spent exploring the object during the test session was considered a measure of object memory in that context. If animals were able to retrieve the original object-context memory representation they should show a decrease in the percentage of object exploration. We found no difference in object exploration time between groups during the TR (Fig. 1C, 1F), but, critically, there was a significant condition effect over the percentages of object exploration during the TS. Animals in the NC condition had significantly higher percentages of object exploration than the AC and PC conditions (Fig. 1D, 1G). Percentages of object exploration were significantly lower than 100% for both the AC and PC, but not for the NC in both versions of the task. The absence of differences due to the cue removal in the PC condition suggests that rats are able to retrieve the original representation using degraded information.

Rearing behavior is considered an alternative measure of environmental novelty, less dependent on object memory than object exploration time [55, 56]. The percentage of rearings of the TS with respect to TR showed a similar tendency as for the percentage of object exploration (Fig. S1).

### Contextual cues in the AR task are important to guide reactivation of the object memory in the Prh

To further strengthen our results, we sought a more direct link between memory retrieval and the reactivation of the hippocampal memory representations. Under particular conditions, memory reactivation can increase susceptibility to protein synthesis inhibitors post-retrieval, which is specific to the memory trace reactivated. Memories that are not active during the test do not become labile and are not sensitive to inhibition of protein synthesis [57-62]. If contextual information is enough to guide retrieval of the original object-in-context memory in our task, exposure to the context should lead to object memory destabilization in regions where this memory is stored, like the perirhinal cortex (Prh) [57, 60, 63]. To address this question, animals trained in the 2-day version of the AR task were exposed to a retrieval session in the same context but in the absence of an object in either AC or NC conditions (TS1). Immediately after context exposure, they were infused into the Prh with the protein synthesis inhibitor Emetine or with Vehicle. Twenty-four hours later, the object memory was evaluated (TS2), by placing the original object next to a novel one (Fig. 2A). Vehicle infused animals had significantly higher discrimination ratios than zero during this last session, evidencing that 10-min of training were enough for a 48h-memory of the object. On the other hand, Emetine infusion in the Prh significantly decreased the discrimination ratio in the AC condition when compared to Vehicle, while no effect was seen in the NC condition (Fig.2B) (One sample t test against 0: NC-Veh *t*=4.77 *p*=.003, NC-Eme *t*=4.22 *p*=.008, AC-Veh *t*=4.38 *p*=.005, AC-Eme *t*=1.30 *p*=.252). These results indicate that the context associated with the object is sufficient to guide the reactivation of the original “object-in-context” memory.

To rule out non-specific effects of Emetine, we tested the contextual dependency of memory labilization. During the training session, rats were exposed to an object in the NC condition and after 2 h to another object in the AC condition. 24h later, animals were exposed to either the empty NC context or the empty AC context and both groups received Emetine or Vehicle infusion immediately after context exposure. Memory for the original objects was tested 24h later against a novel object in both cases (Fig. 2C). Time spent exploring each object during training did not differ between the AC and NC context (Fig. 2D). Nevertheless, a RM Two Way ANOVA indicates a significant effect of drug and type of condition over the discrimination ratios (Fig. 2E). Animals infused with Emetine after NC exposure only showed a reduction of the discrimination ratio for the object originally associated with that condition, while animals infused with Emetine after AC condition had selective reduction for the discrimination of the object associated with AC during the training session. This indicates that the effect of Emetine on memory reconsolidation is specific for the object-context association because only the trace that was previously associated with the presented context was sensitive to Emetine infusion. Additionally, this suggests that all the contexts are capable of guiding reactivation of the relevant object memory (the one associated with each context), while memories of the non-relevant objects are not susceptible to the action of protein synthesis inhibitors.

### CA3 place cells remapping correlates with memory recall

In order to evaluate how context representation is related to episodic memory, we recorded CA3 activity as animals performed the 1-day version of the task (four rats, 27 sessions, n = 332 place cells). We compared place cell firing properties between the TR and TS phases using two parameters that describe similarities and differences between their place fields in both conditions, spatial correlation and firing rate change. We found that a place cell can maintain the position where they fire (similar location of its place field, high spatial correlation) and its firing rate (low firing rate change) or change one or both variables at the same time (Fig 3A) without changing its spatial information coding (Fig. S3A). When we grouped place cells according to the TS condition, we found that the spatial correlation differed significantly between the NC and the other two conditions (Fig. 3B, n=332, Kruskal-Wallis test NC-AC p= .0048, NC-PC p= .0108 and AC-PC p= .8523), but not the firing rate change (Fig. 3C, n=332, Kruskal-Wallis test NC-AC p= .6012, NC-PC p= .1467 and AC-PC p= .4754). Interestingly, even after removing half of the cues (PC), there were no significant differences in the spatial correlation or firing rate change of the place cells when compared with the AC condition (Fig. 3B). This result is in agreement with the behavioral output observed in the PC condition in which behavior was not different between PC and AC (Fig. 1G), and suggests that retrieval of the original contextual memory in the PC context is directly associated with CA3 neuronal representation by place cells.

One advantage of the present task is its variability. We could find sessions in which, despite having reduced the number of cues (NC or PC condition), the animals showed low exploration percentage and sessions where animals exhibit high exploration percentage under a degraded context. This behavioral characteristic was useful to look for a relationship between place cell activity and the memory output (represented as the object exploration percentage) independently of the experimental condition. There was a significant inverse correlation between spatial correlation and the percentage of object exploration and a significant positive correlation between firing rate change and the percentage of object exploration. Thus, animals tend to explore the object more as firing rate change in CA3 increases and spatial correlation decreases (Fig. S3B,C, spatial corr p = 1.47e-10 R = -.34 and firing rate change p =.00015 R = .20).

According to dichotomic accounts of memory retrieval, the behavioral output of the animals should be represented by only two distributions, i.e., retrieval of the previous experience/high place cell activity correlation or no retrieval/low place cell activity correlation. To understand if our behavioral data fitted this dichotomic account of memory, we performed a Bootstrap Likelihood Ratio Test (LRT) for assessing the number of mixture gaussian components that could model our behavioral data. We found that the addition of a second distribution significantly increased the ability of the mixture gaussian component model to explain our data (LRT, p=0.001), but adding a third gaussian component did not lead to a significant improvement (LRT, p=0.997). This analysis suggests that the behavioral output of our animals can be modeled by two independent distributions (i.e., memory retrieval/no memory retrieval). We then classified the recording sessions in terms of the animal’s dichotomic behavioral output (i.e., putative internal context representation) instead of the experimental cue setup (AC, PC or NC). We separated sessions in which animals discriminated between contexts (high exploration) from sessions in which they did not (low exploration), using an estimated 2-SD threshold for context discrimination (see methods, Fig 3D). With this new classification, we found significant differences in the spatial correlation (Fig 3E, n=332, LMM p =1.822e-06) and in the firing rate change (Fig 3F, n=332, LMM p = .0038). To rule out nonspecific effects, we repeated the same analysis but using the activity of the neurons that weren’t classified as place cells and found no differences between groups (Fig. S3D,E). In addition, we fitted our neuronal data with a general linear model (GLM) with a binomial distribution, using the putative internal context representation as the response variable. Models including only one of the variables (spatial correlation or firing rate change) or both were significant (internal rep ∼ spatial corr, p =1.27e-08; internal rep ∼ FR change, p = 1.63e-05 and internal rep ∼ spatial corr + FR change, p spatial corr = 1.97e-06, p FR change = .02). The best of the three models is the one that includes both variables (ANOVA, p 1-variable model vs p 2-variables model = 0.009). From these results we can conclude that place cells are encoding more than only the physical space; instead, their activity is related to the memory that animals express in each context.

### The role of NMDAR during associative retrieval of episodic-like memories

Previous studies have shown that the CA3 region [40, 41], and in particular NMDARs, are required when retrieval occurs in the absence of some of the original contextual cues [45]. These studies used KO models to evidence NMDAR requirements that do not allow for an accurate description of the time window of this requirement. Thus, for this study, we used a pharmacological approach to evaluate the requirement of CA3-NMDARs for retrieval under cue degraded conditions. Animals trained in the 2-day version of the AR task received an injection of the NMDAR antagonist AP5 or Vehicle in the CA3 region 15 min before the test session in the AC or PC condition (Fig. 4A). Vehicle-infused animals spent significantly less time exploring the original object, independently of the condition. However, animals that received an AP5 infusion previous to the test session in the PC condition showed significantly higher exploration percentages than Vehicle infused animals. In contrast, no effect of AP5 was observed in the AC condition (Fig. 4B). Our results suggest that the NMDAR activity in the CA3 region is involved in recovering memories in the presence of partial contextual information but not with all the original cues.

Considering our previous results on the reactivation of the contextual representations under degraded cues, we asked whether NMDARs were required for reactivation of the object-context memory under degraded cues. Consistent with our previous result (Fig 2), Emetine infusion in the Prh immediately after context exposure (TS1) significantly reduced the discrimination ratio of the previously presented object even under a cue degraded context (Fig 4C). But when we combined pre-TS1 infusion of AP5 into the CA3 with post-TS1 infusion of Emetine into Prh, Emetine only reduced the discrimination ratio of the object memory under AC but not under PC context exposure, suggesting that inhibiting NMDAR activity during PC exposure can interfere with the ability to activate the representation of the original memory. The PC specificity of AP5-effect during the exposure session rules out an effect due to changes in memory labilization induced by AP5. Furthermore, the absence of any significant difference in the number of rearings exclude possible AP5-induced changes in exploration levels during the exposure session (paired t test: AC Mean±SEM Veh=36.65±5.84 AP5=42.53±32.77, *t*=0.78, *p*=.444, PC Mean±SEM Veh=56.9±4.37 AP5=53.05±4.89, *t*=0.84, *p*=.409). In sum, these results showed that NMDARs in the CA3 region are not required for memory reactivation when guided by the original contextual information, but are crucial for the reactivation of object-in-context memories under partial contextual information when a memory completion process is required.

### Bidirectional modulation of NMDAR activity in the DG-CA3 circuit affects the balance between memory retrieval and acquisition under cue degraded conditions

Many findings indicate the importance of NMDAR in the HP (in particular the DG) for correct pattern separation function, both at the behavioral [42] and electrophysiological level [43, 64-66]. Interfering with NMDARs could hinder memory differentiation thereby increasing memory generalization. To address this, we tested the role of NMDAR activity in the DG for retrieval under the AC, PC conditions and an additional PC2 condition in which only two cues were left (Fig. 5A). We found no effect of AP5 infusions in the DG 15 min previous to the test session in the AC or PC condition (Fig. 5B). However, using the two-cue context (PC2) during the test session, we observed that DG-Vehicle infused animals showed percentages of exploration that did not differ significantly from training, while AP5 infused animals had a percentage of exploration significantly lower than 100% and significantly different from the Vehicle infused group (Fig. 5C). The lack of effects in less degraded AC and PC conditions rule out any non-specific influence of AP5 over exploration levels. Our results suggest that NMDARs in DG are necessary for the orthogonalization that allows a distinction to be formed between a cue degraded context and the original full context, and that the absence of this distinction could lead to memory retrieval even in highly degraded contexts.

However, we predicted that we could favor memory discrimination by activating NMDAR in the DG. We tested this prediction in animals trained in the 2-day version of the AR task that received an infusion of the NMDAR partial agonist D-Cycloserine (D-Cyc) or Vehicle in the DG 15 min before exposure to a PC context (Fig. 5D). Animals infused with D-Cyc in the DG before TS showed significantly higher exploration percentages than the Veh group, with object exploration times that did not differ from TR (Fig. 5E). These results indicate that NMDAR activity in the DG can decrease the ability to generalize from a degraded context. In contrast, we already showed that NMDARs in the CA3 region are important for contextual generalization. We reasoned that we could increase retrieval guided by incomplete cue stimuli through an increase in NMDAR activity in the CA3. We used a similar strategy than the one for the DG by presenting a PC2 context that was not sufficient for an effective object in context memory retrieval (Fig. 5D and F). We infused D-Cyc or Vehicle before the test phase under the PC2 context and found that Veh-infused animals showed exploration levels consistent with a lack of object in context memory retrieval. However, the infusion of D-Cyc in the CA3 previous to this test phase significantly decreased the percentage of object exploration (Fig.5F), indicating that increasing NMDAR activity in the CA3 region can enhance memory retrieval under cue degraded conditions, potentially through tipping the balance towards contextual generalization.

Changes in engram cell excitability [67] or plasticity [12, 13, 68] could be controlling the efficacy of memory retrieval under contextually degraded conditions in our task. In this regard, NMDA receptor activation can trigger AMPA-type trafficking to the dendritic surface in an L-type voltage gated calcium channel (L-VGCC)-dependent manner [69] and L-VGCC can also regulate neuronal excitability [70, 71]. Therefore, we evaluated their role as potential effectors of NMDARs. Animals were infused in CA3 with the L-type calcium channel inhibitor Nimodipine (Nim) previous to the D-Cyc infusion and then tested in the PC2 context (Fig.5G). While D-Cyc infusion decreased the percentage of object exploration, co-infusion of D-Cyc with Nimodipine prevented this effect. This result suggests that NMDARs interact with L-VGCC in the CA3 region to favor memory generalization. Histological analysis did not reveal any lesion in the infusion sites (CA3 or DG) (Supplementary Fig.2). Results cannot be explained by drug-related changes in motivation to explore the object during the test phase, since infusion of D-Cyc in the DG or CA3 region 15 min before exposure to an object in the AC context did not alter exploration percentages compared with Vehicle-infused groups (Paired t test: *t*_DG_=0.67*p*=.528 n=7; *W*_CA3_=-8 *p*=.578, n=7). In summary, our results suggest that by bidirectional manipulation of NMDARs in the CA3/DG region affects the hippocampal processing balance between a retrieval mode (favored by generalization like processing) and an acquisition mode (favored by a differentiation like processing).

We performed a dichotomic analysis to account for how these drugs putatively affect memory retrieval, similar to what we used for electrophysiological experiments (i.e., putative internal context representation). We divided the data in two groups by pooling together all “memory boosting” (Left) or “memory impairing” (Right) drug infusion experiments (see methods, Fig. 6A). We found that the “memory boosting” group, in a two-cue context where most animals show a behavior non consistent with retrieval, the treatment increased the proportion of rats that behaved in a retrieval-like fashion (has a similar internal context representation between TS and TR, Fig. 6B left). In turn, in the “memory impairing” group, under a three-cue context where most animals behave in a retrieval-like fashion, the treatment decreased the number of rats that behaved in a manner consistent with retrieval (Fig. 6B right). These results show that, although there is a great variability in animal behavior, NMDAR activity in the hippocampal circuit can bias the internal spatial representation towards memory retrieval or away from it.

**Fig. 6.**
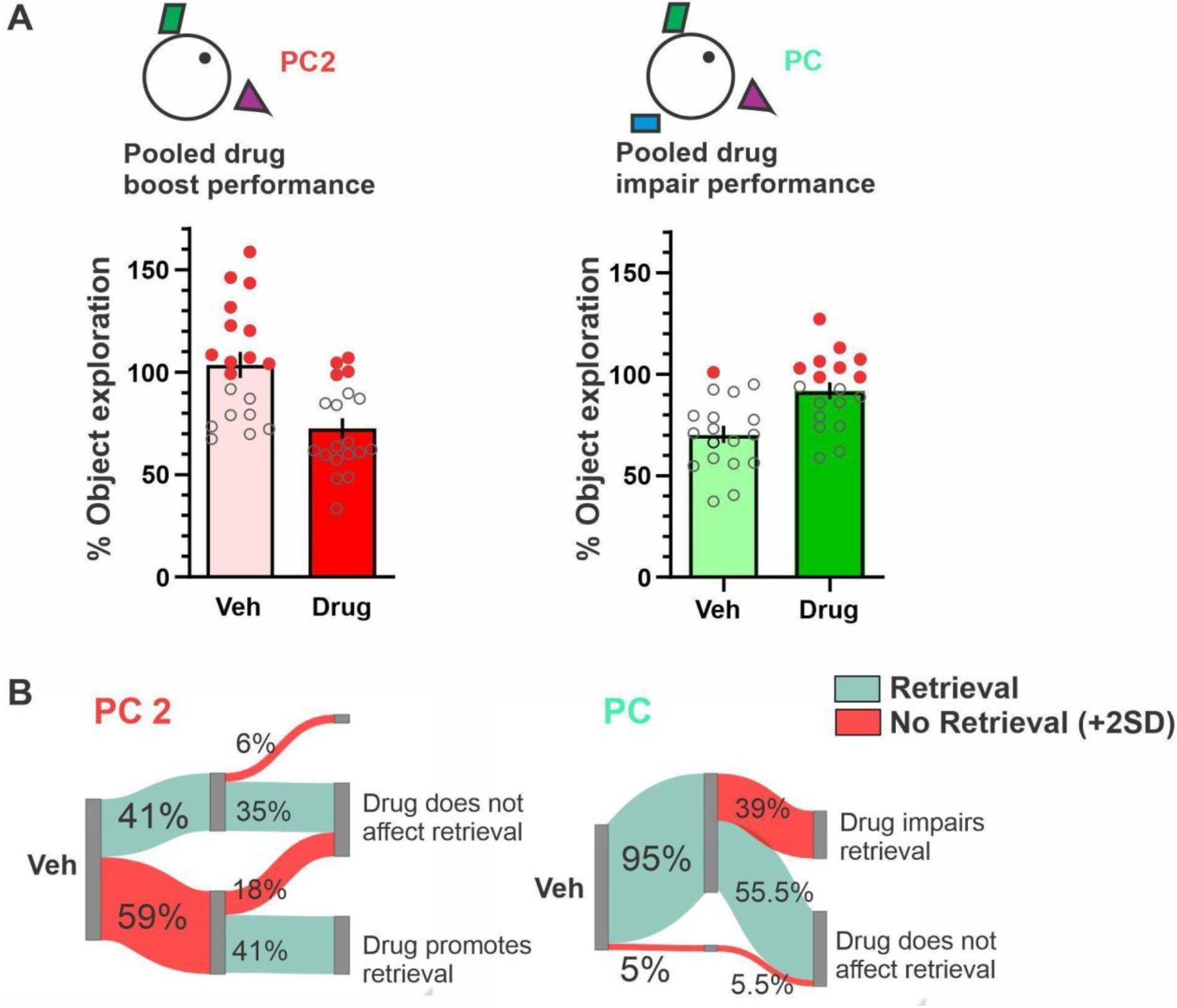
Drugs affect animals’ context representation in a cue-degraded environment. **(A)** Pooled distribution of all “memory boosting” (Left) or “memory impairing” (Right) drug infusion experiments, depicting as red dots the animals above the memory retrieval threshold (i.e. internal context representation, 2SD threshold taken from independent values of Fig.1 B AC-PC). **(B)** Percentage of animals that were above 2SD retrieval threshold (red) during Veh session or below retrieval threshold (green), showing how much of each percentage is affected by the drug or remains unchanged. (?2=3.125 p=0.03855, pool drug impair performance, ?2=5.14286 p=0.0167, one tail McNemar chiSquare test)

## Discussion

Memory is a flexible process that adapts to the organisms’ requirements in a changing environment. The conditions for retrieval of memories that are not specifically associated with reward or punishment are likely more dependent on both the cues present in the environment and the internal state of the individual. In this study we investigated the molecular and neural mechanisms involved in the recovery of spatial representations stored as incidental memories of an object experienced in a particular context. Our findings indicate that memory differentiation and memory generalization functions compete for behavioral control. We showed that the amount of remapping of CA3 neurons, that signals a separation process, is related to the recall of an episodic-like object-in-context memory. When animals recognized an object in context as familiar, regardless of the contextual cues, there was less remapping in CA3 neurons than when animals recognized the relationship as novel. This finding is crucial for understanding how these contextual representations can influence episodic memory recall and behavior. In the same line, we showed that NMDAR activity in the DG-CA3 circuit can influence this discrimination/generalization balance. We found that pharmacological treatments that favor pattern separation (Fig. 4B, 5E) tend to make animals behave as if the relationship between object and context is novel while treatments that promote pattern completion (Fig. 5C, F) bias animals to retrieve a previous experience.

Holistic retrieval of memories is considered a key element in episodic memory allowing the recovery of all aspects of an event in an integrated manner. Our results indicate that, even if distal cues provide essential allocentric information for spatial navigation, animals could retrieve a memory of an object in a particular context, guided by a limited amount of contextual information. This is consistent with other studies in rodents [40, 45] and humans [72]. Besides, the similar patterns of object exploration and rearings in the presence of a variable number of retrieval cues indicates that exploration levels of familiar objects in the present task can be directly related to contextual novelty.

Place cells in the hippocampus can remap in response to contextual changes [73], enabling different activity patterns to represent different environments [74, 75]. But none of those previous studies showed if there is any behavioral meaning in remapping (i.e. is there any link between remapping and memory?). There are interesting studies showing that artificial activation of a selected group of hippocampal neurons can modify mice behavior in a contextual fear memory [76-79] or a reward-oriented paradigm [80]. Here we go a step further and show that the degree of spontaneous incidental memory, that does not involve any reward or punishment, correlates with the amount of remapping of CA3 place cells. When animals recognize an environment as unfamiliar there is more remapping than when they experience the context as familiar. This result is fundamental for a better understanding of contextual representations and their role for memory, without the influence of explicit motivational factors.

Our results evidence a large variability between (and within) animals in terms of how these contextual representations, coded by place cell activity, change with contextual variation. This has been repeatedly shown in the literature [32, 81, 82]. Even more, place cells in CA1 could remap within the same session even though there are no changes in the context, possibly due to modifications in the internal state of the animal [32]. This variability could be explained if remapping does not solely reflect the observable and controlled properties of the environment, but rather subjective perceptions and predictions the animal is making about the environment during retrieval [83]. However, the variability observed could also be due to individual variability in the encoding of the environment between trials that results in the animal being able to retrieve the original memory in some trials but not in others. In fact, interindividual and intraindividual variability in behavior is more the rule than the exception in behavioral studies and it only recently has caught the attention of researchers [84-87]. However, the relationship between remapping of these representations and the behavioral output of memory recall remained unclear. Interestingly, we showed that the amount of memory retrieved during TS in the AR task was not only reliant on the TS condition itself (number of cues) but was specifically related to the animal’s internal representation of the context. So, by dividing sessions only by TS condition, we were missing interesting data regarding the behavioral outputs. Instead, sorting remapping by animal’s internal representation reflects more clearly the different encoding of distinct contexts for the animal (Fig 3.D-F). This contrast between the effect of the contextual manipulation (AC, PC, NC) and the actual contextual representational change reveals that the controlled experimental manipulations of the context only reflect a subset of what the animal perceives as the context. Accordingly, the individual representation of the same space is variable and can be more or less subject to discrimination or generalization depending on uncontrolled experimental variables that increase their importance as the context becomes more degraded. This relationship became evident when we tried to explain object exploration (as a measure of memory retrieval) by spatial correlation and firing rate change. The level of object exploration was better explained by both variables taken together, suggesting that both spatial correlation and firing rate change are essential for memory discrimination. In addition, the effect of the pharmacological manipulations over the generalization/discrimination was also variable and likely dependent on the retrieval state of the animal (Fig. 6B). We propose that the pharmacological treatments biased the distribution of exploration times towards discrimination or generalization changing the proportion of individuals performing pattern separation or completion that would depend on their internal representation activation in the TS.

In the labilization/reconsolidation experiments (Fig. 2E, 4C), both contextual and object information are neither degraded nor absent in the PC condition. Thus, object memory retrieval demands a reconstruction of the original experience from degraded information to guide memory. Combined, our results show that incidental reactivation occurs for all the elements of the original event even in contextually degraded conditions, and that retrieval is related to a low degree of remapping in CA3 place cells. In addition, these results reveal a contextual specificity of the reactivation and subsequent labilization of object memory traces, since only the corresponding contextually associated object memory trace was reactivated under contextual exposure in the NC and AC retrieval sessions (Fig.2E). Consistent with this idea, human studies found that activity in the Prh correlates with recollection in response to partial spatial cues [88, 89] and associative retrieval led to neocortical activity corresponding to incidental reactivation of all the items of a particular event [72].

Our results are consistent with other studies that suggest a requirement of NMDAR activity in the CA3 region for long term spatial memory reactivation, but only when the amount of contextual information available is limited [45, 90]. Since spatial place fields are generated and modified in an NMDAR-dependent manner [64-66, 91-94], it is possible that memory reactivation in cue-degraded conditions (where presumably the type of information entry over the perforant path is reduced) relies more heavily on the activity of attractor states and the strength of recurrent collateral connections in CA3. In this context, changes in NMDAR activity could affect the attractor circuit activation threshold since enhancement of CA3 recurrent collateral connections is NMDAR-dependent [10, 12, 13, 95]. Therefore, NMDAR antagonists could be preventing reverberant activity in the CA3 region, leading to a retrieval deficit, and NMDAR agonists could favor attractor circuit stability for the corresponding memory trace, thereby promoting retrieval. Could memory retrieval under contextually degraded cues rely on LTP expression mechanisms that involve an NMDAR-dependent component in CA3? Currently, it seems counterintuitive to think of a role of NMDARs in the expression of associative LTP, because, for many years, the vast majority of research focused on the role of NMDAR for LTP induction and not expression [96, 97]. Nonetheless, a recent publication showed that recall cues can drive transient increases in excitability specific to engram cells and this in turn can influence the accuracy and efficacy of memory retrieval [67]. Interestingly, synaptic activation of NMDARs triggers internalization of Kir2.1 channels leading to increases in engram cell excitability. Partial cue driven NMDAR-dependent increases in cell excitability in the CA3 region could be controlling the efficacy of memory retrieval under contextually degraded conditions in our task. This is consistent with previous results that indicate that the absence of NMDARs affects both spatial memory and stable place field activity [45] and with several lines of evidence suggesting that stabilization and posterior reactivation of place fields require NMDAR activity and protein synthesis [66, 98-100]. The fact that a voltage gated calcium channel blocker (Nimodipine) prevented the increase in memory generalization due to CA3 D-Cyc infusion points toward a role of calcium channels and possible excitability changes regulating the strength of cue-driven retrieval under partial cue conditions.

Computational models suggest that the attractor circuit in CA3 could lead to pattern separation or pattern completion, as a function of the relative strength of the attractor circuit and the nature of the external inputs from the EC and the DG [5, 6, 101]. Some models predict that sensory information entering through the perforant path should either be used to recover the original engram in CA3 and cause retrieval of the entire memory, or to form a new engram guided by the new entrance of information. Thereby, these theories imply that recognition of an experience will ultimately depend on whether pattern completion-like processes or pattern separation-like processes rule over CA3 activation patterns. This is exactly what we have seen in the electrophysiological data, in which the degree of remapping balances between retrieval of a familiar memory with low remapping or recognizing a different situation with no retrieval (and accordingly higher remapping). In this context, we speculate that, under AC and PC conditions, input from the EC through the perforant path could lead to attractor activity in the CA3 region corresponding to an object in context representation and, by the time DG inputs arrive, the strong attractor dynamics of the circuit could dominate over the DG inputs and prevent remapping favoring a pattern completion process. But if the sensory inputs induced by retrieval cues are weaker, like in the PC2 condition, CA3 attractor circuit activity will also be weaker, offering the opportunity for these attractor states to follow DG input and promote environmentally induced remapping.

In this scenario, we have shown that interfering with plasticity related mechanisms in the DG can actually favor memory retrieval in conditions in which spatial cues would normally be insufficient to guide retrieval. It is possible that inactivation of NMDAR, that is known to affect rate remapping [42] should alter the ability of the DG to remap in response to novel stimuli, leading to stable memory representation in the CA3 region and memory retrieval under incomplete contextual information. Electrophysiological studies that support this line of thought have shown that LTP decay is an active process that requires NMDAR activity [102]. Consistent with the putative DG role in pattern separation, NMDAR partial agonist D-Cyc prevented memory retrieval in the PC condition, suggesting a shift in the hippocampal balance towards an acquisition mode instead of a retrieval mode. This is in line with the result showing that granular cells of the DG are depolarized after the animal has been exposed to a novel environment, indicating that these cells could be involved in setting the hippocampal circuit into an acquisition mode [103]. Moreover, although most studies focused on the role of NMDARs on spatial memory acquisition [104-107], there are a few studies that already showed a post-acquisition facilitation by administration of NMDAR antagonists [102, 108].

Although in real life, episodic memory retrieval usually occurs in a degraded context, this process and the balance between pattern completion - pattern separation that it entails have not been studied under those conditions. In addition, there has been relatively little work done linking remapping and behavior (see *Allegra et al*. 2020) Using an incidental memory task, we showed that retrieval of a context memory is reflected by the levels of CA3 remapping, demonstrating the relationship between remapping an episodic-like memory. Furthermore, we describe NMDARs as a key player in the regulation of the balance between retrieval and memory differentiation processes. While increase in CA3 NMDAR activity boosts memory retrieval, DG NMDAR activity enhances the memory differentiation process. Our results contribute to understanding the adaptive nature of memory to guide behavior consistent with changes in the environmental cues and the internal state of the individual.

## Acknowledgements

We thank Bárbara Giugovaz Tropper and Victoria Deschamps for her technical assistance, David Jaime for his help with animal care and maintenance, and Dr.Jorge Medina for helpful discussions and sharing of some of the drugs. This work was supported by CONICET (PUE 0052) to PB, FONCYT (PICT 2018-1062) to PB and CONICET (PIP 2656) to MB.

## Supplementary Material

**Fig. S1.**
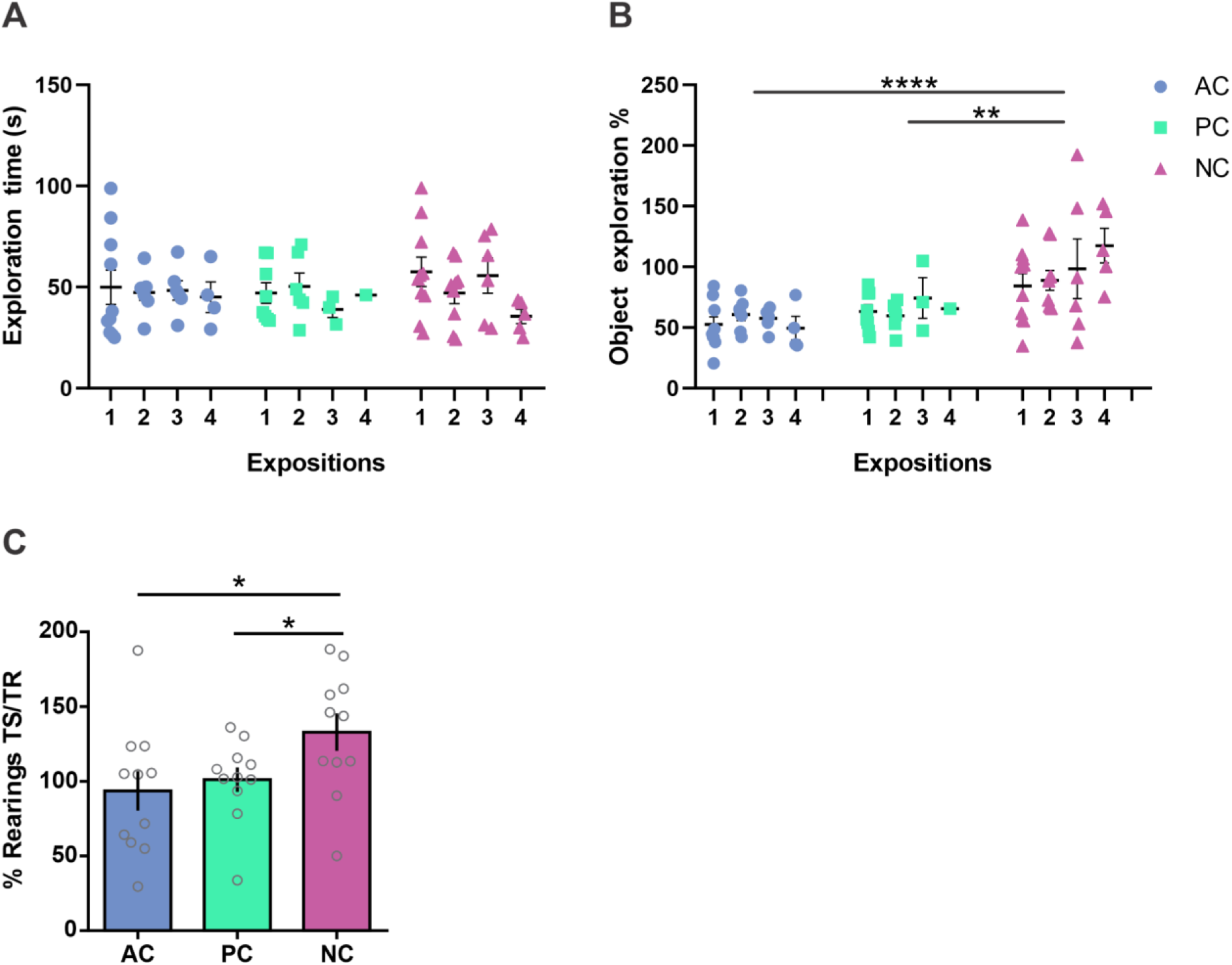
**(A) and (B)** Repeated measure analysis. **(A)** Total object exploration during the training session for AC, PC and NC in the different expositions to the same condition during the 1-day version of the task. Two way RM ANOVA, n=10, sessions = 75, each dot represents a session, interaction: F =.5755, p = .7484 **(B)** Total object exploration percentage for AC, PC and NC in the different expositions to the same condition during the 1-day version of the task. Two way RM ANOVA, n=10, sessions = 75, each dot represents a session, F = 16.04, p < .0001.Tukey’s post hoc test: **** p<.0001 AC vs NC; p=.25 AC vs PC ; ** p =.001 NC vs PC. One sample t test against 100% AC t =14.45, p<.0001; PC t=8.76, p<.0001; NC t = 0.37 p=.91. The re-exposures to the different conditions have no effect on the exploration time in the training session or in the object exploration percentage. **(C)** Percentage of rearings during the test phase in relation to training for the AC, PC and NC conditions in the 2-day version. One way RM ANOVA F=4.26, p=.048, n=9. * p <.05, ** p <.01. While no previous differences between groups in rearings were present during the training session (RM One Way ANOVA, F=0.30, p=0.69), post-hoc comparisons revealed a marginally significant increase in the NC condition when compared to both the AC and PC conditions (tAC/NC=2.23, pAC/NC=.05, tPC/NC=2.14, pPC/NC=.06). To note, unlike object-related exploration, contextual exploration in the AC condition (as manifested by the number of rearings) does not decrease between TR and TS since animals were habituated to the AC context (and not to the object) previously to TR session (One sample t test from 100%, AC t=0.49, p=.636, PC t=0.13, p=0.896, NC t=2.62, p=.025). Individual values used to calculate the mean and SEM are presented as dots.

**Fig S2.**
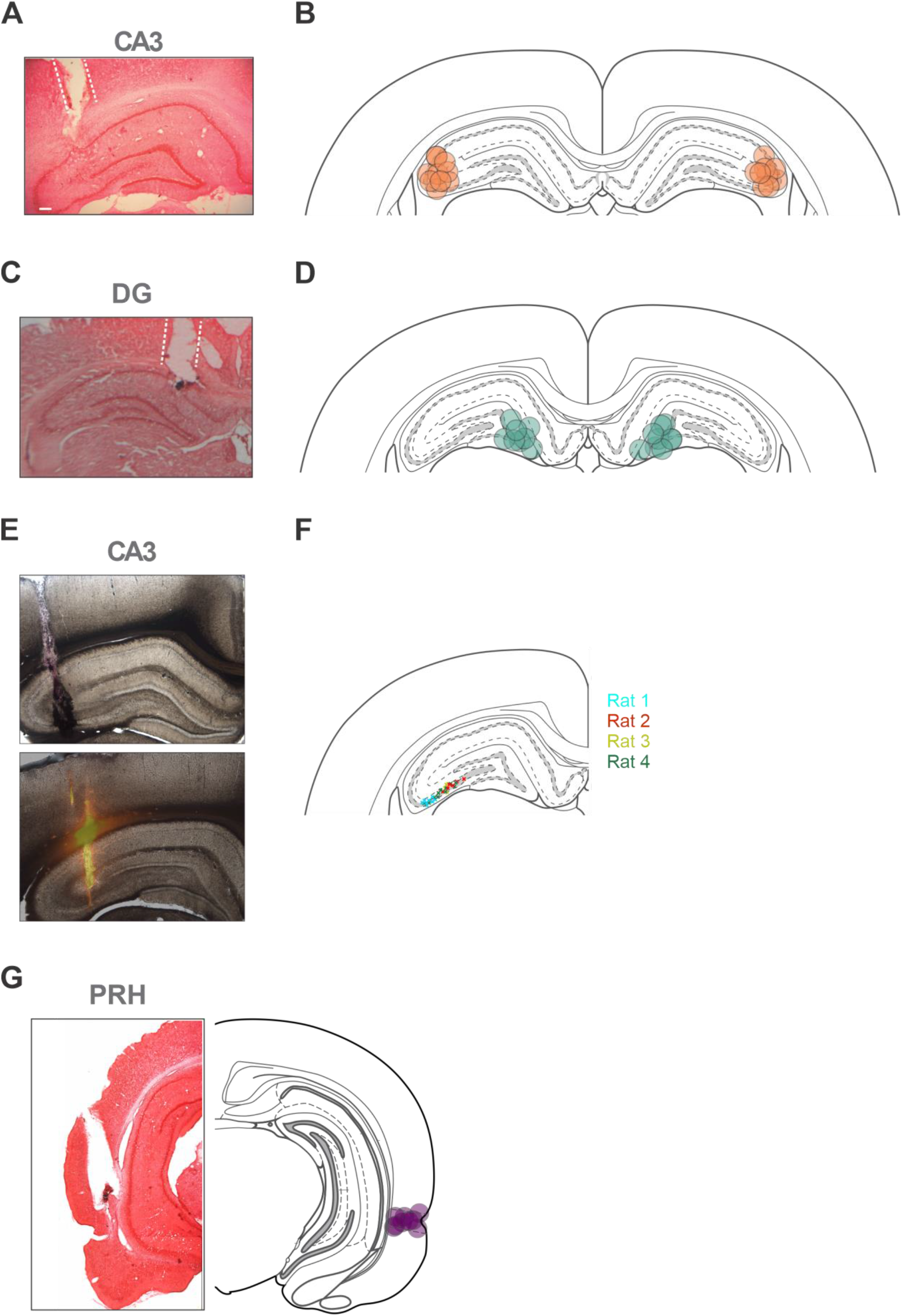
**(A)** Safranin staining image of a coronal brain section of a rat cannulated in the CA3 of HP. **(B)** Schematic representation of coronal sections of the rat brain in the rostrocaudal plane of -3.60 taken from Atlas Paxinos and Watson. Circles represent a representative infusion area in an example experiment. **(C)** Safranin staining image of a coronal brain section of a rat cannulated in the CA3 of HP. **(D)** Schematic representation of coronal sections of the rat brain in the rostrocaudal plane of - 3.90 taken from Atlas Paxinos and Watson. Circles represent representative infusion areas of example experiment. **(E)** Two representative coronal brain sections with tetrode trayectory. **(F)** Tetrode recording sites for 3 recorded rats in CA3. Each cross represents a tetrode tip. See full methods for verification of tetrode recording sites. **(G)** Safranin staining image of a coronal brain section of a rat cannulated in the PRH. Scale 200 μm.

**Fig S3.**
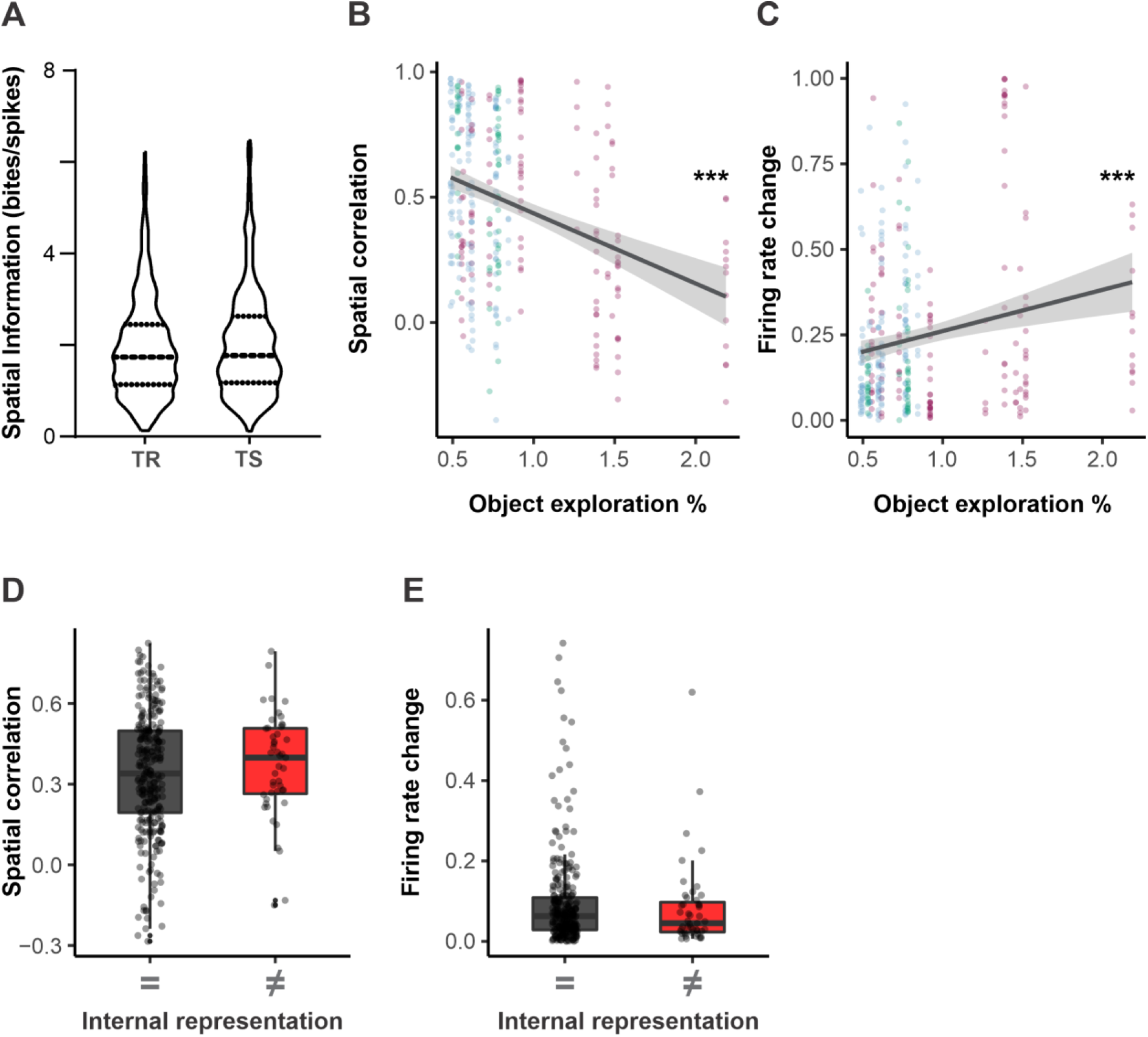
**(A)** No change in spatial coding betweenTR and TS. n = 332 Paired T-test p = .14 **(B) and (C)** Correlation between place cell activity and memory output. **Spatial correlation:** Pearson’s correlation between spatial correlation and object exploration %, n=332, p = 1.47e-10 R = -.34. **Firing rate change:** Pearson’s correlation between firing rate change and object exploration %, n=332, p =.00015 R = .20 **(D) and (E)** No place cells don’t encode animal internal representation of context. The same number of neurons, 332, that in figure 2 were selected randomly from the subset of record neurons that didn’t fulfill the place cell criteria and the same analysis that in figure 2 was applied. **(D)** No place cell **spatial correlation** sorted by internal representation. n=332 Wilcoxon rank-sum test, p = .39 **(E)** No place cell **firing rate change** sorted by internal representation. n=332 Wilcoxon rank-sum test, p= .36

## Bibliography

1. Rumelhart, D.E., J.L. McClelland, and University of California San Diego. PDP Research Group., Parallel distributed processing : explorations in the microstructure of cognition. Computational models of cognition and perception. 1986, Cambridge, Mass.: MIT Press.

2. McClelland, J.L., B.L. McNaughton, and R.C. O’Reilly, Why there are complementary learning systems in the hippocampus and neocortex: insights from the successes and failures of connectionist models of learning and memory. Psychol Rev, 1995. 102(3): p. 419–57.

3. Norman, K.A. and R.C. O’Reilly, Modeling hippocampal and neocortical contributions to recognition memory: a complementary-learning-systems approach. Psychol Rev, 2003. 110(4): p. 611–46.

4. Kohonen, T., Associative memory : a system-theoretical approach. Communication and cybernetics v 17. 1977, Berlin ; New York: Springer-Verlag. ix, 176 p.

5. Marr, D., Simple memory: a theory for archicortex. Philos Trans R Soc Lond B Biol Sci, 1971. 262(841): p. 23–81.

6. Rolls, E.T., et al., Information about spatial view in an ensemble of primate hippocampal cells. J Neurophysiol, 1998. 79(4): p. 1797–813.

7. O’Reilly, R.C. and J.W. Rudy, Computational principles of learning in the neocortex and hippocampus. Hippocampus, 2000. 10(4): p. 389–97.

8. Amaral, D.G. and M.P. Witter, The three-dimensional organization of the hippocampal formation: a review of anatomical data. Neuroscience, 1989. 31(3): p. 571–91.

9. O’Reilly, R.C. and J.L. McClelland, Hippocampal conjunctive encoding, storage, and recall: avoiding a trade-off. Hippocampus, 1994. 4(6): p. 661–82.

10. Harris, E.W. and C.W. Cotman, Long-term potentiation of guinea pig mossy fiber responses is not blocked by N-methyl D-aspartate antagonists. Neurosci Lett, 1986. 70(1): p. 132–7.

11. Zalutsky, R.A. and R.A. Nicoll, Comparison of two forms of long-term potentiation in single hippocampal neurons. Science, 1990. 248(4963): p. 1619–24.

12. Ito, I., et al., Synapse-selective impairment of NMDA receptor functions in mice lacking NMDA receptor epsilon 1 or epsilon 2 subunit. J Physiol, 1997. 500 (Pt 2): p. 401–8.

13. Debanne, D., B.H. Gahwiler, and S.M. Thompson, Long-term synaptic plasticity between pairs of individual CA3 pyramidal cells in rat hippocampal slice cultures. J Physiol, 1998. 507 (Pt 1): p. 237–47.

14. O’Keefe, J. and J. Dostrovsky, The hippocampus as a spatial map. Preliminary evidence from unit activity in the freely-moving rat. Brain Res, 1971. 34(1): p. 171–5.

15. Pastalkova, E., et al., Internally generated cell assembly sequences in the rat hippocampus. Science, 2008. 321(i5894): p. 1322–7.

16. MacDonald, C.J., et al., Hippocampal “time cells” bridge the gap in memory for discontiguous events. Neuron, 2011. 71(4): p. 737–49.

17. Sakurai, Y., Hippocampal and neocortical cell assemblies encode memory processes for different types of stimuli in the rat. J Neurosci, 1996. 16(8): p. 2809–19.

18. Wood, E.R., P.A. Dudchenko, and H. Eichenbaum, The global record of memory in hippocampal neuronal activity. Nature, 1999. 397(6720): p. 613–6.

19. Eichenbaum, H., et al., Cue-sampling and goal-approach correlates of hippocampal unit activity in rats performing an odor-discrimination task. J Neurosci, 1987. 7(3): p. 716–32.

20. Komorowski, R.W., J.R. Manns, and H. Eichenbaum, Robust conjunctive item-place coding by hippocampal neurons parallels learning what happens where. J Neurosci, 2009. 29(31): p. 9918–29.

21. Manns, J.R., M.W. Howard, and H. Eichenbaum, Gradual changes in hippocampal activity support remembering the order of events. Neuron, 2007. 56(3): p. 530–40.

22. Terada, S., et al., Temporal and Rate Coding for Discrete Event Sequences in the Hippocampus. Neuron, 2017. 94(6): p. 1248–1262 e4.

23. Leutgeb, S., et al., Distinct ensemble codes in hippocampal areas CA3 and CA1. Science, 2004. 305(5688): p. 1295–8.

24. Guzowski, J.F., J.J. Knierim, and E.I. Moser, Ensemble dynamics of hippocampal regions CA3 and CA1. Neuron, 2004. 44(4): p. 581–4.

25. Neunuebel, J.P. and J.J. Knierim, CA3 retrieves coherent representations from degraded input: direct evidence for CA3 pattern completion and dentate gyrus pattern separation. Neuron, 2014. 81(2): p. 416–27.

26. O’Keefe, J., Place units in the hippocampus of the freely moving rat. Exp Neurol, 1976. 51(1): p. 78–109.

27. Tolman, E.C., Cognitive maps in rats and men. Psychol Rev, 1948. 55(4): p. 189–208.

28. Eichenbaum, H., et al., The hippocampus, memory, and place cells: is it spatial memory or a memory space? Neuron, 1999. 23(2): p. 209–26.

29. O’keefe, J.N. L., The Hippocampus as a Cognitive Map. Oxford University Press, 1978.

30. Leutgeb, S., et al., Place cells, spatial maps and the population code for memory. Curr Opin Neurobiol, 2005. 15(6): p. 738–46.

31. Gauthier, J.L. and D.W. Tank, A Dedicated Population for Reward Coding in the Hippocampus. Neuron, 2018. 99(1): p. 179–193 e7.

32. Pettit, N.L., X.C. Yuan, and C.D. Harvey, Hippocampal place codes are gated by behavioral engagement. Nat Neurosci, 2022. 25(5): p. 561–566.

33. Kentros, C.G., et al., Increased attention to spatial context increases both place field stability and spatial memory. Neuron, 2004. 42(2): p. 283–95.

34. Law, L.M., D.A. Bulkin, and D.M. Smith, Slow stabilization of concurrently acquired hippocampal context representations. Hippocampus, 2016. 26(12): p. 1560–1569.

35. Yonelinas, A.P., et al., Recollection and familiarity: examining controversial assumptions and new directions. Hippocampus, 2010. 20(11): p. 1178–94.

36. Squire, L.R., J.T. Wixted, and R.E. Clark, Recognition memory and the medial temporal lobe: a new perspective. Nat Rev Neurosci, 2007. 8(11): p. 872–83.

37. Rolls, E.T., A computational theory of episodic memory formation in the hippocampus. Behav Brain Res, 2010. 215(2): p. 180–96.

38. Leutgeb, J.K., et al., Pattern separation in the dentate gyrus and CA3 of the hippocampus. Science, 2007. 315(5814): p. 961–6.

39. Gilbert, P.E., R.P. Kesner, and I. Lee, Dissociating hippocampal subregions: double dissociation between dentate gyrus and CA1. Hippocampus, 2001. 11(6): p. 626–36.

40. Gold, A.E. and R.P. Kesner, The role of the CA3 subregion of the dorsal hippocampus in spatial pattern completion in the rat. Hippocampus, 2005. 15(6): p. 808–14.

41. Kesner, R.P., M.R. Hunsaker, and M.W. Warthen, The CA3 subregion of the hippocampus is critical for episodic memory processing by means of relational encoding in rats. Behav Neurosci, 2008. 122(6): p. 1217–25.

42. McHugh, T.J., et al., Dentate gyrus NMDA receptors mediate rapid pattern separation in the hippocampal network. Science, 2007. 317(5834): p. 94–9.

43. Nakazawa, K., et al., Hippocampal CA3 NMDA receptors are crucial for memory acquisition of one-time experience. Neuron, 2003. 38(2): p. 305–15.

44. Rolls, E.T., A quantitative theory of the functions of the hippocampal CA3 network in memory. Front Cell Neurosci, 2013. 7: p. 98.

45. Nakazawa, K., et al., Requirement for hippocampal CA3 NMDA receptors in associative memory recall. Science, 2002. 297(5579): p. 211–8.

46. Paxinos, G. and C. Watson, The rat brain in stereotaxic coordinates. 4th ed. 1998, San Diego: Academic Press.

47. Myers, R.D., Injection of solutions into cerebral tissue: Relation between volume and diffusion. Physiol Behav, 1966. 1(2): p. 171–174.

48. Lomax, P., The distribution of morphine following intracerebral microinjection. Experientia, 1966. 22(4): p. 249–50.

49. Hazan, L., M. Zugaro, and G. Buzsaki, Klusters, NeuroScope, NDManager: a free software suite for neurophysiological data processing and visualization. J Neurosci Methods, 2006. 155(2): p. 207–16.

50. Harris, K.D., et al., Accuracy of tetrode spike separation as determined by simultaneous intracellular and extracellular measurements. J Neurophysiol, 2000. 84(1): p. 401–14.

51. Markus, E.J., et al., Spatial information content and reliability of hippocampal CA1 neurons: effects of visual input. Hippocampus, 1994. 4(4): p. 410–21.

52. Ennaceur, A. and J. Delacour, A new one-trial test for neurobiological studies of memory in rats. 1: Behavioral data. Behav Brain Res, 1988. 31(1): p. 47–59.

53. Winters, B.D., L.M. Saksida, and T.J. Bussey, Object recognition memory: neurobiological mechanisms of encoding, consolidation and retrieval. Neurosci Biobehav Rev, 2008. 32(5): p. 1055–70.

54. Blaser, R. and C. Heyser, Spontaneous object recognition: a promising approach to the comparative study of memory. Front Behav Neurosci, 2015. 9: p. 183.

55. Sturman, O., P.L. Germain, and J. Bohacek, Exploratory rearing: a context- and stress-sensitive behavior recorded in the open-field test. Stress, 2018. 21(5): p. 443–452.

56. Sarafino, E., Experiential aspects of exploratory behavior in rats. 1978: p. 235–243.

57. Morici, J.F., et al., 5-HT2a receptor in mPFC influences context-guided reconsolidation of object memory in perirhinal cortex. Elife, 2018. 7.

58. Radiske, A., et al., BDNF controls object recognition memory reconsolidation. Neurobiol Learn Mem, 2017. 142(Pt A): p. 79–84.

59. Winters, B.D., et al., On the dynamic nature of the engram: evidence for circuit-level reorganization of object memory traces following reactivation. J Neurosci, 2011. 31(48): p. 17719–28.

60. Rossato, J.I., et al., On the role of hippocampal protein synthesis in the consolidation and reconsolidation of object recognition memory. Learn Mem, 2007. 14(1): p. 36–46.

61. Nader, K., Memory traces unbound. Trends Neurosci, 2003. 26(2): p. 65–72.

62. Alberini, C.M., The role of reconsolidation and the dynamic process of long-term memory formation and storage. Front Behav Neurosci, 2011. 5: p. 12.

63. Dudai, Y. and M. Eisenberg, Rites of passage of the engram: reconsolidation and the lingering consolidation hypothesis. Neuron, 2004. 44(1): p. 93–100.

64. Dupret, D., et al., The reorganization and reactivation of hippocampal maps predict spatial memory performance. Nat Neurosci, 2010. 13(8): p. 995–1002.

65. Hayashi, Y., NMDA Receptor-Dependent Dynamics of Hippocampal Place Cell Ensembles. J Neurosci, 2019. 39(26): p. 5173–5182.

66. Kentros, C., et al., Abolition of long-term stability of new hippocampal place cell maps by NMDA receptor blockade. Science, 1998. 280(5372): p. 2121–6.

67. Pignatelli, M., et al., Engram Cell Excitability State Determines the Efficacy of Memory Retrieval. Neuron, 2019. 101(2): p. 274–284 e5.

68. Williams, J.M., et al., Differential trafficking of AMPA and NMDA receptors during long-term potentiation in awake adult animals. J Neurosci, 2007. 27(51): p. 14171–8.

69. Hiester, B.G., et al., L-Type Voltage-Gated Ca(2+) Channels Regulate Synaptic-Activity-Triggered Recycling Endosome Fusion in Neuronal Dendrites. Cell Rep, 2017. 21(8): p. 2134–2146.

70. Gamelli, A.E., et al., Deletion of the L-type calcium channel Ca(V) 1.3 but not Ca(V) 1.2 results in a diminished sAHP in mouse CA1 pyramidal neurons. Hippocampus, 2011. 21(2): p. 133–41.

71. Moore, S.J. and G.G. Murphy, The role of L-type calcium channels in neuronal excitability and aging. Neurobiol Learn Mem, 2020. 173: p. 107230.

72. Horner, A.J., et al., Evidence for holistic episodic recollection via hippocampal pattern completion. Nat Commun, 2015. 6: p. 7462.

73. Muller, R.U. and J.L. Kubie, The effects of changes in the environment on the spatial firing of hippocampal complex-spike cells. J Neurosci, 1987. 7(7): p. 1951–68.

74. Leutgeb, S., et al., Independent codes for spatial and episodic memory in hippocampal neuronal ensembles. Science, 2005. 309(5734): p. 619–23.

75. Leutgeb, S., et al., Fast rate coding in hippocampal CA3 cell ensembles. Hippocampus, 2006. 16(9): p. 765–74.

76. Cowansage, K.K., et al., Direct reactivation of a coherent neocortical memory of context. Neuron, 2014. 84(2): p. 432–41.

77. Redondo, R.L., et al., Bidirectional switch of the valence associated with a hippocampal contextual memory engram. Nature, 2014. 513(7518): p. 426–30.

78. Liu, X., et al., Optogenetic stimulation of a hippocampal engram activates fear memory recall. Nature, 2012. 484(7394): p. 381–5.

79. Ramirez, S., et al., Creating a false memory in the hippocampus. Science, 2013. 341(6144): p. 387–91.

80. Robinson, N.T.M., et al., Targeted Activation of Hippocampal Place Cells Drives Memory-Guided Spatial Behavior. Cell, 2020. 183(6): p. 1586–1599 e10.

81. Fenton, A.A. and R.U. Muller, Place cell discharge is extremely variable during individual passes of the rat through the firing field. Proc Natl Acad Sci U S A, 1998. 95(6): p. 3182–7.

82. Kelemen, E. and A.A. Fenton, Coordinating different representations in the hippocampus. Neurobiol Learn Mem, 2016. 129: p. 50–9.

83. Sanders, H., M.A. Wilson, and S.J. Gershman, Hippocampal remapping as hidden state inference. Elife, 2020. 9.

84. Kastner, D.B., et al., Spatial preferences account for inter-animal variability during the continual learning of a dynamic cognitive task. Cell Rep, 2022. 39(3): p. 110708.

85. Koh, M.T., R.W. McMahan, and M. Gallagher, Individual differences in neurocognitive aging in outbred male and female long-evans rats. Behav Neurosci, 2022. 136(1): p. 13–18.

86. Pittaras, E., H. Hamelin, and S. Granon, Inter-Individual Differences in Cognitive Tasks: Focusing on the Shaping of Decision-Making Strategies. Front Behav Neurosci, 2022. 16: p. 818746.

87. Belkaid, M., et al., Mice adaptively generate choice variability in a deterministic task. Commun Biol, 2020. 3(1): p. 34.

88. Diana, R.A., A.P. Yonelinas, and C. Ranganath, Imaging recollection and familiarity in the medial temporal lobe: a three-component model. Trends Cogn Sci, 2007. 11(9): p. 379–86.

89. Sakai, K. and Y. Miyashita, Neural organization for the long-term memory of paired associates. Nature, 1991. 354(6349): p. 152–5.

90. Fellini, L., et al., Pharmacological intervention of hippocampal CA3 NMDA receptors impairs acquisition and long-term memory retrieval of spatial pattern completion task. Learn Mem, 2009. 16(6): p. 387–94.

91. Cabral, H.O., et al., Single-trial properties of place cells in control and CA1 NMDA receptor subunit 1-KO mice. J Neurosci, 2014. 34(48): p. 15861–9.

92. McHugh, T.J., et al., Impaired hippocampal representation of space in CA1-specific NMDAR1 knockout mice. Cell, 1996. 87(7): p. 1339–49.

93. Tonegawa, S., et al., Hippocampal CA1-region-restricted knockout of NMDAR1 gene disrupts synaptic plasticity, place fields, and spatial learning. Cold Spring Harb Symp Quant Biol, 1996. 61: p. 225–38.

94. Tsien, J.Z., P.T. Huerta, and S. Tonegawa, The essential role of hippocampal CA1 NMDA receptor-dependent synaptic plasticity in spatial memory. Cell, 1996. 87(7): p. 1327–38.

95. Nicoll, R.A., J.A. Kauer, and R.C. Malenka, The current excitement in long-term potentiation. Neuron, 1988. 1(2): p. 97–103.

96. Muller, D., M. Joly, and G. Lynch, Contributions of quisqualate and NMDA receptors to the induction and expression of LTP. Science, 1988. 242(4886): p. 1694–7.

97. Bawin, S.M., W.M. Satmary, and W.R. Adey, Roles of the NMDA and quisqualate/kainate receptors in the induction and expression of kindled bursts in rat hippocampal slices. Epilepsy Res, 1993. 15(1): p. 7–13.

98. Agnihotri, N.T., et al., The long-term stability of new hippocampal place fields requires new protein synthesis. Proc Natl Acad Sci U S A, 2004. 101(10): p. 3656–61.

99. Ekstrom, A.D., et al., NMDA receptor antagonism blocks experience-dependent expansion of hippocampal “place fields”. Neuron, 2001. 31(4): p. 631–8.

100. Dragoi, G., K.D. Harris, and G. Buzsaki, Place representation within hippocampal networks is modified by long-term potentiation. Neuron, 2003. 39(5): p. 843–53.

101. McNaughton, B.L. and R.G. Morris, Hippocampal synaptic enhancement and information storage within a distributed memory system. Trends Neurosci, 1987. 10(10).

102. Villarreal, D.M., et al., NMDA receptor antagonists sustain LTP and spatial memory: active processes mediate LTP decay. Nat Neurosci, 2002. 5(1): p. 48–52.

103. Gomez-Ocadiz, R., et al., A synaptic signal for novelty processing in the hippocampus. Nat Commun, 2022. 13(1): p. 4122.

104. Bannerman, D.M., J.N. Rawlins, and M.A. Good, The drugs don’t work-or do they? Pharmacological and transgenic studies of the contribution of NMDA and GluR-A-containing AMPA receptors to hippocampal-dependent memory. Psychopharmacology (Berl), 2006. 188(4): p. 552–66.

105. Bast, T., B.M. da Silva, and R.G. Morris, Distinct contributions of hippocampal NMDA and AMPA receptors to encoding and retrieval of one-trial place memory. J Neurosci, 2005. 25(25): p. 5845–56.

106. Morris, R.G., et al., N-methyl-d-aspartate receptors, learning and memory: chronic intraventricular infusion of the NMDA receptor antagonist d-AP5 interacts directly with the neural mechanisms of spatial learning. Eur J Neurosci, 2013. 37(5): p. 700–17.

107. Morris, R.G., et al., Selective impairment of learning and blockade of long-term potentiation by an N-methyl-D-aspartate receptor antagonist, AP5. Nature, 1986. 319(6056): p. 774–6.

108. Shinohara, K. and T. Hata, Post-acquisition hippocampal NMDA receptor blockade sustains retention of spatial reference memory in Morris water maze. Behav Brain Res, 2014. 259: p. 261–7.

109. Allegra, M., et al., Differential Relation between Neuronal and Behavioral Discrimination during Hippocampal Memory Encoding. Neuron, 2020. 108(6): p. 1103–1112 e6.

